# The International Space Station selects for microorganisms adapted to the extreme environment, but does not induce genomic and physiological changes relevant for human health

**DOI:** 10.1101/533752

**Authors:** Maximilian Mora, Lisa Wink, Ines Kögler, Alexander Mahnert, Petra Rettberg, Petra Schwendner, René Demets, Charles Cockell, Tatiana Alekhova, Andreas Klingl, Alina Alexandrova, Christine Moissl-Eichinger

**Affiliations:** Medical University of Graz, Department of Internal Medicine, Graz, Austria; German Aerospace Center (DLR), Institute of Aerospace Medicine, Radiation Biology Department, Research Group Astrobiology, Cologne, Germany; University of Edinburgh, School of Physics and Astronomy, Edinburgh, UK; European Space Research and Technology Centre (ESTEC), Noordwijk, The Netherlands; Lomonosov Moscow State University, Biological Faculty, Moscow, Russia; Ludwig Maximilians University of Munich, Plant Development and Electron Microscopy, Department of Biology I, Biocenter, Planegg-Martinsried, Germany; BioTechMed Graz, Austria

**Keywords:** International Space Station, ISS, microbiome, antibiotics resistance

## Abstract

The International Space Station (ISS) is a unique, completely confined habitat for the human crew and co-inhabiting microorganisms. Here, we report on the results of the ISS experiment “EXTREMOPHILES”. We aimed to exploit the microbial information obtained from three surface and air sampling events aboard the International Space Station during increments 51 and 52 (2017) with respect to: i) microbial sources, diversity and distribution within the ISS, ii) functional capacity of microbiome and microbial isolates, iii) extremotolerance and antibiotics-resistance (compared to ground controls), and iv) microbial behavior towards ISS-relevant materials such as biofilm formation, or potential for degradation. We used wipe samples and analyzed them by amplicon and metagenomics sequencing, cultivation, comparative physiological studies, antibiotic resistance tests, genome analysis of isolates and co-incubation experiments with ISS-relevant materials. The major findings were: i) the ISS microbiome profile is highly similar to ground-based confined indoor environments, ii) the ISS microbiome is subject to fluctuations and indicative for the (functional) location, although a core microbiome was present over time and independent from location, iii) the ISS selects for microorganisms adapted to the extreme environment, but does not necessarily induce genomic and physiological changes which might be relevant for human health, iv) cleanrooms and cargo seems to be a minor source of microbial contamination aboard, and v) microorganisms can attach to and grow on ISS-relevant materials. Biofilm formation might be a threat for spacecraft materials with the potential to induce instrument malfunctioning with consequences for mission success. We conclude that our data do not raise direct reason for concern with respect to crew health, but indicate a potential threat towards biofilm formation and material integrity in moist areas.

## Introduction

Human space exploration beyond boundaries of Earth and Moon is a declared goal of NASA, ESA, Roscosmos and other space-faring agencies, envisaging a potential human Mars mission in the next 20 to 30 years. Maintenance of astronauts’ health during a several hundred days journey in a confined artificial environment in space is one of the key aspects which has to be addressed for such a long-term mission.

The human immune system was shown to be compromised under space flight conditions, as a significant decrease of lymphocytes and also of the activity of innate and adaptive immune response was observed when compared to terrestrial controls (1,2). Adding an order of complexity, human health is strongly intertwined with its microbiome, billions of microorganisms thriving on external and internal surfaces of the human body.

Our body’s microbiome is dependent on the environmental microbiome, as they are in constant exchange and interaction. Isolated human subjects in hospitals were observed to loose microbial diversity, serving as an indicator for health and stability in general (Koskinen 2019, unpublished); this might, however, be contradictory to observations on the astronauts’ gut microbiome (3). The microbiome is prone to external factors, which influence the composition or the function of the microbial community. For instance, microbial diversity in built environments was shown to negatively correlate with the diversity of antimicrobial resistances especially in confined habitats in a recent publication (Mahnert et al, 2019, unpublished). With respect to space travel, it has been shown that microgravity affects the virulence of certain microorganisms, such as *Salmonella typhimurium* (4), *Listeria monocytogenes* and *Enterococcus faecalis* (5). As a consequence, monitoring of the microbial community aboard spacecraft is highly important to assess risk factors to the health of crew members.

Additionally, some microorganisms might even pose a risk to the material integrity of a spacecraft: So-called technophilic microorganisms, in particular fungi, are able to corrode alloys and polymers used in spacecraft assembly (6). Technophilic microorganisms caused major problems on the former Russian space station Mir (7,8).

The majority of information with respect to environmental microbiome dynamics aboard manned spacecraft is retrieved from ground-based simulation studies, such as the Mars500 (9) and the HI-SEAS (http://hi-seas.org/) experiments. However, the ISS is currently representing the most isolated human habitat.

The ISS circles our planet in low Earth orbit (approx. 400 km above ground) and is constantly inhabited since more than 18 years. Except for cargo exchange and the arrival of new crew members roughly every six months, the ISS is completely sealed off from any surrounding biological ecosystem and thus represents one of the most isolated and confined man-made environments to date (10). Like no other currently available testbed for long-term manned space missions, the ISS has the scientific benefit of providing real spaceflight conditions, including microgravity and an elevated background radiation - parameters, which are hard to implement in ground-based simulation studies.

The ISS consists of different modules, and while new modules were added over the years, also the crew size increased. Nowadays six international astronauts and cosmonauts routinely inhabit the ISS, along with their associated microorganisms. While certain microorganisms might benefit from the constant temperature (approx. 22°C) and stable humidity (approx. 60%) aboard the ISS, aforementioned harsh spaceflight conditions as well as low nutrient levels due to regular cleaning and a reduced introduction of new material, make the ISS for microorganisms a unique and extreme-situated indoor environment (11).

Recent publications focused on the microbial analysis of ISS debris and dust (12–15), the study of the astronauts’ microbiome (16), the characterization of bacterial and fungal isolates from the ISS (17,18) and the (molecular) microbial analysis of swab and wipe samples taken inside the ISS (19). A study investigating the growth behavior of non-pathogenic (terrestrial) bacteria aboard the ISS found no changes in most bacteria, given that they have enough nutrients (11).

Other publications also focused on the detection of antimicrobial resistance genes aboard the ISS and evaluated the potential risk these genes might represent in a closed spacecraft environment (20). Singh et al. assessed the succession and persistence of microbial communities and the associated antimicrobial resistance and virulence properties based on metagenomic reads obtained from samples of three flights. Overall, 46 microbial species were found, including eight biorisk group 2 species, to be persistent on the ISS over a timespan of roughly one and a half years (21). The authors inferred an increase of antimicrobial resistance and virulence genes over time bearing an alarming message that these factors seem to become an increasing proportion of the ISS microbiome.

Although an increased risk for the health of the astronauts and cosmonauts aboard has been proposed several times based on the molecular detection of antimicrobial resistances, viruses and pathogens, infections of crew members or health issues related to pathogenic action of microorganisms have been reported only rarely (22). Moreover, the given environmental microbial contamination limits (air, surfaces) were exceeded only in a few cases to date, in which appropriate countermeasures succeeded in a timely manner (23). Moreover, a recent genomics-based study could not reveal potentially health-threatening differences in *Bacillus* and *Staphylococcus* pangenomes from ISS, compared to human-associated and soil pangenomes (24).

In this study, we report on the realization of the ISS experiment EXTREMOPHILES, targeting the profile, diversity, dynamics and functional capacity of the microbiome aboard. We used cultivation efforts to obtain microbial isolates. We assessed their genomic and physiological adaptation towards ISS conditions and tested the hypothesis, whether, as indicated by previous literature reports, ISS microorganisms possess a higher extremo-tolerance and antibiotics-resistance potential compared toground controls. Moreover, we were interested in the surface-microbe interaction with regard to material integrity, exhibited by selected, freshly-isolated ISS strains.

## Materials and Methods

### Pre-flight preparations and sampling aboard the ISS

Packaging, pre-processing and logistics of the sample material regarding upload and download from the ISS were managed by the Biotechnology Space Support Center (BIOTESC) of the Lucerne University of Applied Sciences and Arts (Switzerland). In-flight sampling aboard the ISS was performed during increment 51 and 52 (April to June 2017) under the ESA (European Space Agency) operation named EXTREMOPHILES. Sampling was performed either with dry wipes (session A and B) or pre-moistened wipes (session C; TX 3211 Alpha Wipe, ITW Texwipe, Kernersville, US, 23×23 cm; 20 ml autoclaved ultra-pure water for chromatography, LiChrosolv, Merck Millipore). Three sampling sessions were performed, with session A on 01.05.2017, session B on 12.07.2017 and session C on 29.06.2017. With a span of 72 days between session A and B they were conceptualized for comparative sampling to assess microbiome fluctuation over time. An overview of all sampled areas and sessions is given in Table 1.

**Table 1:**
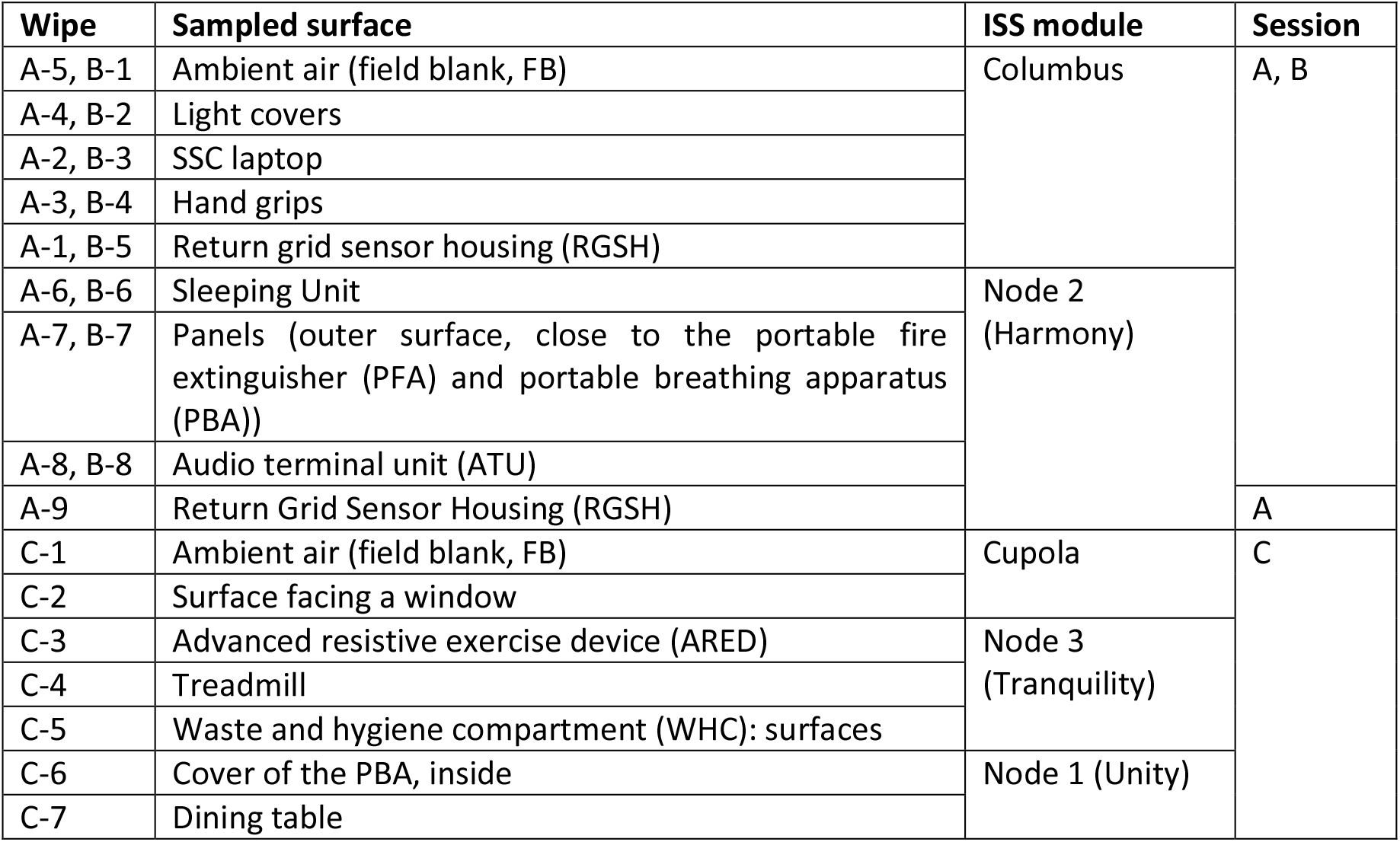
Sampling locations and sampling sessions. In total, 24 wipes were retrieved from 5 different modules, and 15 different locations within the ISS.

The sampling instructions for each session were as follows. 1: Put on sterile gloves (DNA-free nitrile gloves, ABF Diagnostics GmbH, Kranzberg, Germany). 2: Using gloved hand, remove wipe X from bag (metal closure bag, GML-alfaplast GmbH, Munich, Germany), wave wipe through the air (approx. 20s). Put wipe back into its bag and close properly. 3: Change glove. 4: Using gloved hand, take sample using wipe Y according to Table 1. Put wipe back into its bag and close properly. 5: Repeat steps 3 and 4 for every new sampling surface according to Table 1. 6: Store wipes at ambient (session A, B, dry wipes), or under cool conditions (“cold stowage”, 2-10°C, session C, moist wipes).

### Cleanroom and cargo vehicle sampling

In order to retrieve samples for comparative analyses, one ISS-relevant cleanroom and cargo-spacecraft was sampled, namely cleanroom S5C at the Centre Spatial Guayanais near Kourou in French Guiana, housing ATV5 “Georges Lemaître”. Swab (FLOQSwabs™, Copan diagnostics, USA) and wipe samples from ATV and its cleanroom were provided by Stefanie Raffestin (ESA) in 2014.

### Sample extraction

The obtained sample material was either available as wipes or swabs (cleanroom). Wipes were submerged in 80 ml DNA-free 0.9% (w/v) NaCl solution (NaCl was heat-treated to destroy DNA residues for 24 hours, 250°C), vortexed (10s) and shaken manually (15s), ultra-sonicated at 40 kHz for 2 min and vortexed (10s). The sampling material was aseptically removed from the extraction solution before cultivation- and molecular analyses. Swabs were submerged in 15 ml of NaCl solution and processed identically.

### Cultivation

Cultivation of microorganisms was performed on a number of solid and liquid media, as given in Table 2. For microbial enrichment, we provided variable chemical and physical conditions with respect to: pH (pH 4-10), temperature (4-65°C), gas phase (aerobic, N_2:_H_2:_CO_2_, H_2:_CO_2_, N_2:_CO_2_), nutrients and nutrient availability. R2A (pH5-7), RAVAN and ROGOSA were supplemented with nystatin (50 μg/ml) to suppress growth of fungi; media targeting archaea were supplemented with 50 μg/ml streptomycin and 100 μg/ml ampicillin. Inoculation was done using 500 and 250 μl (duplicates) of the extraction solution. In addition, 500 μl aliquots of the wipe suspension were irradiated at the DLR in Cologne, Germany, to select for radiation resistant isolates. They were either irradiated by UV-C (254nm) with an intensity of 50 J/m^2^, 75 J/m^2^, 100 J/m^2^, and 200 J/m^2^ or by X-Rays with an intensity of 125 Gy, 250 Gy, 500 Gy, 750 Gy, or 1000 Gy. Radiation resistant microorganisms were cultivated on R2A and TSA agar. Pure cultures were obtained via repeated dilution series in liquid medium and/or purification streaks on solid media.

**Table 2:** Cultivation conditions and media.

| Medium | Abb. | Target organisms | T [°C] | pH | Phase | Gas phase | Reference or manufacturer |
| --- | --- | --- | --- | --- | --- | --- | --- |
| R2A pH 4 | R4 | Acidophiles | 32 | 4 | Solid | Aerobic (ambient) | VWR Chemicals<br>BDH Prolabo® |
| R2A pH 5 | R5 | Acidotolerants | 32 | 5 | Solid | Aerobic (ambient) |  |
| R2A pH 7 | R7W | Mesophiles | 32 | 7 | Solid | Aerobic (ambient) |  |
|  | R7K | Psychrophiles | 4 | 7 | Solid | Aerobic (ambient) |  |
|  | R7A | Anaerobes | 32 | 7 | Solid | N<sub>2</sub>:H<sub>2</sub>:CO<sub>2</sub> (80:10:10) |  |
| R2A pH 9 | R9 | Alkalitolerants | 32 | 9 | Solid | Aerobic (ambient) |  |
| R2A pH 10 | R10 | Alkaliphiles | 32 | 10 | Solid | Aerobic (ambient) |  |
| RAVAN | RAV | Oligotrophs | 32 | 7 | Solid | Aerobic (ambient) | (25) (1:100 modified) |
| Halo medium | HLO | Halophiles | 40 | 7.5 | Solid | Aerobic (ambient) | DSMZ medium 97<br>(https://www.dsmz.de/) |
| Rogosa | ROG | Lactobacilli | 32 | 5.5 | Solid | Aerobic (ambient) | Merck |
| Tryptic Soy Agar | TSA | Mesophiles | 32 |  | Solid | Aerobic (ambient) | Becton Dickinson |
| Yeast-extract peptone medium | YPD | Yeasts & Fungi | 32 | 7 | Solid | Aerobic (ambient) | Sigma-Aldrich |
| Potato dextrose agar | PDA | Yeasts & Fungi | 25 | 5.6 | Solid | Aerobic (ambient) | Sigma-Aldrich |
| Autotrophic homoacetogen medium | AHM | Homoacetogen | 32 | 7 | Liquid | H<sub>2</sub>:CO<sub>2</sub> (20:80) | (26) |
| Autotrophic allrounder medium | AAM | Autotrophs | 32 | 7 | Liquid | N<sub>2</sub>:CO<sub>2</sub> (80:20) | (26) |
| Archaea supporting medium | ASM | Archaea | 32 | 7 | Liquid | H<sub>2</sub>:CO<sub>2</sub> (20:80) | (26) |
| Medium for methanogens | MS | Methanogens | 40 | 7 | Liquid | H<sub>2</sub>:CO<sub>2</sub> (20:80) | (27) |
| MS+organics | MSO | Methanogens | 40 | 7 | Liquid | H<sub>2</sub>:CO<sub>2</sub> (20:80) | This study |
| *N. exaquare* medium | NEX | Thaumarchaea | 32 | 8.5 | Liquid | Aerobic (ambient) | (28) |
| Mixed ruminal bacteria medium | MCB-3 | Methanogens | 40 | 7 | Liquid | H<sub>2</sub>:CO<sub>2</sub> (20:80) | DSMZ medium322<br>(https://www.dsmz.de/) |
| R2A pH 7 liquid | R2A | Thermophiles | 65 | 7 | Liquid | Aerobic (ambient) | This study |

### 16S rRNA gene sequencing and classification of the bacterial and fungal isolates

Partial 16S rRNA genes of the isolates were amplified using the primers 9bF (5’-GRGTTTGATCCTGGCTCAG-3’) and 1406uR (5’-ACGGGCGGTGTGTRCAA-3’), applying the following cycling conditions: Initial denaturation at 95°C for 2min, followed by 10 cycles of denaturing at 96°C for 30s, annealing at 60°C for 30s and elongation at 72°C for 60s, followed by another 22 cycles of denaturing at 94°C for 30s, annealing at 60°C for 30s and elongation at 72°C for 60s, and a final elongation step at 72°C for 10min (29). The template was either a small fraction of a picked colony in a colony-PCR assay or 5–20ng of DNA purified from culture via the peqGOLD Bacterial DNA Kit (Peqlab, Germany). The 16S rRNA gene amplicons were visualized on a 1.5% agarose gel, purified with the Min Elute PCR Purification Kit (Qiagen, Netherlands) or the Monarch PCR and DNA Cleanup Kit (New England Biolabs, US). After Sanger-sequencing (Eurofins, Germany) the obtained sequences were classified using the EzBioCloud platform at http://www.ezbiocloud.net/eztaxon (30).

The ITS region of fungal isolates was sequenced using the primers ITS1F (5’-CTTGGTCATTTAGAGGAAGTAA-3’) and ITS4 (5’-TCCTCCGCTTATTGATATGC-3’) and following cycling conditions: initial denaturation at 95°C for 10min, followed by 35 cycles of denaturing at 94°C for 60s, annealing at 51°C for 60s, elongation at 72°C for 60s, and a final elongation step at 72°C for 8min. The amplicons were Sanger-sequenced (Eurofins, Germany) and the obtained sequence was classified using the curated databases UNITE version 7.2 (31) and BOLD version 4 (32). Fungal isolates of session A, B, and C were classified according to phenotypical characteristics

### Phylogenetic tree reconstruction

For phylogenetic tree reconstruction, the forward and reverse sequences obtained from the isolates were merged to reach a minimum sequence length of 1000 bp. The phylogenetic tree was calculated with the Fast Tree programme (33) and displayed with the Interactive Tree of Life online tool iTOL (34).

### DNA extraction of ISS wipe samples

After aliquots were removed for cultivation assays, the rest of the wipe solutions were filled into Amicon Ultra-15 filter tubes (Sigma Aldrich) and were centrifuged at 4000x g for 10-30 min at 4°C. The flow-through was discarded and the remaining liquid in the filters was pipetted into 1.5ml Eppendorf tubes for DNA extraction with the modified XS-buffer method as previously described (35). DNA concentrations were determined using Qubit (Life Technologies, US).

### Microbial profiling using next-generation sequencing methods

To investigate the detectable molecular diversity, we used a “universal” and an Archaea-targeting approach. The 16S rRNA gene amplicons for the universal approach were amplified using Illumina-tagged primers F515 (5’-TCGTCGG-CAGCGTCAGATGTGTATAAGAGACAGGTGCCAGCMGCCGCGGTAA-3’) and R806 (5’-GTCTCGTGGGCTCGGAGATGTGTATAAGAGACAGGGAC-TACHVGGGTWTCTAA3′) (36). Archaeal amplicons were obtained by a nested approach (37): First, a ~550 bp-long 16S rRNA gene amplicon was generated with the primers Arch344F (5’-ACGGGGYGCAGCAGGCGCGA-3’) and Arch915R (5’-GTGCTCCCCCGCCAATTCCT-3’) (38,39) and in a second PCR, the amplicons for Illumina sequencing were generated by the tagged primers Arch519F (5’-TCGTCGGCAGCGTCAGATGTGTATAAGAGACAGCAGCMGCCGCGGTAA-3’) and Arch785R (5’-GTCTCGTGGGCTCGGAGATGTGTATAAGAGACAGGACTACHVGGGTATCTAATCC-3’) (40), using the purified product of the first PCR as template. The cycling conditions for the universal approach were initial denaturation at 94°C for 3 min, followed by 35 cycles of denaturing at 94°C for 45s, annealing at 60°C for 60s and elongation at 72°C for 90s, followed by a final elongation step at 72°C for 10 min. For the first PCR of the nested archaeal approach, the cycling conditions were initial denaturation at 95°C for 2 min, followed by 10 cycles of denaturing at 96°C for 30s, annealing at 60°C for 30s, and elongation at 72°C for 60s, followed by another 15 cycles of denaturing at 94°C for 30s, annealing at 60°C for 30s, and elongation at 72°C for 60s, and a final elongation step at 72°C for 10min. For the second amplification the cycling conditions were initial denaturation at 95°C for 5 min, followed by 25 cycles of denaturing at 95°C for 40s, annealing at 63°C for 120s and elongation at 72°C for 60s, followed by a final elongation step at 72°C for 10 min.

### Genome sequencing, genome reconstruction and annotation of selected isolates

We sequenced the genomic DNA of six isolates obtained from ISS samples described earlier (41). DNA was isolated from overnight cultures using the peqGOLD bacterial DNA mini kit (Peqlab, Germany). Double stranded DNA was quantified via Qubit Fluorometer 2.0 (Invitrogen, USA) according to manufacturer’s instructions. Library preparation and sequencing was carried out at the Core Facility Molecular Biology at the Center for Medical Research at the Medical University Graz, Austria.

Genomic reads were quality checked with FastQC (42) and then filtered with Trimmomatic (removed all adapter sequences, SLIDINGWINDOW 4:20, MINLEN 50)(43). Genomes were assembled with SPADES in careful mode (44) and afterwards checked for completeness via CheckM (45). The assemblies were annotated and compared to closely related reference strains via the microbial genome annotation & analysis platform MicroScope (http://www.genoscope.cns.fr/agc/microscope) (46–48).

### Resistance and physiological tests

Experiments were performed with selected microbial isolates from this and our recent study on ISS microorganisms (14). (i) **Heat-shock resistance test**: The heat-shock test was carried out according to ESA standards (49). In brief, single colonies of 3–5-day old cultures were suspended in two test tubes containing 2.5 ml sterile PBS. One tube was incubated at room temperature (control), whereas the other was placed in a water bath and exposed for 15 min to 80°C. Samples were immediately cooled down on ice for 5 min after incubation time. The temperature was monitored using a separate pilot tube containing 2.5 ml PBS. Afterwards, 0.5 ml of the heat-shocked suspension and 0.5 ml of the room temperature suspension were plated and incubated at 32°C for 72h. (ii) **Physiological tests**: For the assessment of the temperature range, cultures were plated on R2A pH7 agar and incubated overnight at 32°C. Then the incubation temperatures for the species still growing were stepwise decreased and increased until no further growth was observed. Limits of pH tolerance were assessed accordingly. (iii) **Antibiotics susceptibility tests**: Antimicrobial susceptibility testing for selected, clinically relevant antibiotics (Table 3) was performed using Etest^®^ reagent strips (Biomérieux, Germany) according to manufacturer’s instruction and detailed in (14). Since there were no species-specific breakpoints available, MICs were interpreted according to EUCAST guideline table “PK/PD (Non-species related) breakpoints” (50). In brief, overnight cultures (2–3-day cultures for slower-growing bacteria) were suspended in 0.9% saline. 100 μl of this suspension was plated on standardized Müller-Hinton agar for antimicrobial susceptibility testing (Becton Dickinson, USA). Etest^®^ reagent strips were placed on the plates followed by aerobic incubation for 24 h at 34°C.

**Table 3:** Antibiotics used for antimicrobial susceptibility tests, see also (14).

| Antibiotic substance | Type | Mechanism of action/target | Target group | Concentrations applied (µg/ml) |
| --- | --- | --- | --- | --- |
| Amoxicillin/clavulanic acid | β- Lactam antibiotic (penicillin) and β- lactamase inhibitor | Inhibits cell wall synthesis; bactericidal | Gram + /- | 0.016- 256 |
| Ampicillin | β- Lactam antibiotic (penicillin) | Inhibits cell wall synthesis; bactericidal | Gram + /- | 0.016- 256 |
| Cefotaxime | β- Lactam antibiotic (cephalosporin) | Inhibits cell wall synthesis; bactericidal | Gram + /- | 0.016– 32 |
| Ceftriaxone | β- Lactam antibiotic; (cephalosporin) | Inhibits cell wall synthesis; bactericidal | Gram + /- | 0.016– 256 |
| Ciprofloxacin | Fluoroquinolone | Inhibits bacterial DNA gyrase; bactericidal | Gram + /- | 0.002– 32 |
| Clarithromycin | Macrolide | Inhibits protein synthesis; bacteriostatic | Gram + /- | 0.016– 256 |
| Clindamycin | Lincosamide | Inhibits protein synthesis; bacteriostatic | Gram + /- | 0.016– 256 |
| Colistin | Polypeptide antibiotic; polymyxin | Attacks cell membrane; bactericidal | Gram - | 0.016– 256 |
| Doxycycline | Polyketide antibiotic; (tetracycline) | Inhibits protein synthesis; bacteriostatic | Gram + /- | 0.016– 256 |
| Gentamicin | Aminoglycoside | Inhibits protein synthesis; bactericidal | Gram + /- | 0.016– 256 |
| Levofloxacin | Fluoroquinolone | Inhibits bacterial DNA gyrase; bactericidal | Gram + /- | 0.002– 32 |
| Linezolid | Oxazolidinone | Inhibits protein synthesis; bacteriostatic | Gram + | 0.016– 256 |
| Meropenem | β- Lactam antibiotic (carbapenem) | Inhibits cell wall synthesis; bactericidal | Gram + /- | 0.002– 32 |
| Moxifloxacin | Fluoroquinolone | Inhibits bacterial DNA gyrase; bactericidal | Gram + /- | 0.002– 32 |
| Penicillin G | β- Lactam antibiotic (penicillin) | Inhibits cell wall synthesis; bactericidal | Gram+ | 0.016– 256 |
| Trimethoprim/sul famethoxazole | Dihydrofolate reductase inhibitor and sulfonamide | Inhibits tetrahydrofolate synthesis; bactericidal | Gram + /- | 0.002– 32 |
| Vancomycin | Glycopeptide antibiotic | Inhibits cell wall synthesis; bactericidal | Gram+ | 0.016– 256 |

### Co-incubation experiments and electron microscopy

To test if some of our isolates interact with, and possibly damage, materials used aboard the ISS, we incubated them together with relevant ISS material. Pieces of NOMEX^®^ fabric were provided by the Biotechnology Space Support Center (BIOTESC) of the Lucerne University and plates of the aluminum copper magnesium alloy EN AW 2219 which is also used on the ISS, were provided by Thales Alenia Space (TAS), Italy. NOMEX^®^ is a flexible, flameproof fabric used for most storage bags aboard the ISS. The NOMEX^®^ fabric was cut into pieces of 20 mm x 30 mm and autoclaved before incubation. The aluminum alloy EN AW 2219 was cut into small plates of 20 mm x 30 mm x 3 mm by Josef Baumann in Falkenberg, Germany, and then evenly polished with a grit size of P240 and partly eloxated by Heuberger Eloxal, Austria. The autoclaved metal platelets, non-eloxated and eloxated, and NOMEX^®^ fabric pieces were then incubated together with bacteria isolated from the ISS: *Cupriavidus metallidurans* pH5_R2_1_II_A (aerobic), *Bacillus licheniformis* R2A_5R_0.5 (aerobic), and *Cutibacterium avidum* R7A_A1_IIIA (anaerobic). Incubations were done in triplicates over a period of 3 months in liquid R2A medium in Hungate tubes at pH7 and 32°C. Every 2 weeks, 50% of the medium was exchanged to ensure survival and further growth of the bacteria. After incubation, metal plates and NOMEX^®^ fabric pieces were investigated via scanning electron microscopy. Metal plates and NOMEX^®^ fabric pieces from the co-incubation experiment with selected bacteria were aseptically removed from their respective Hungate tube, carefully rinsed with 1xPBS buffer and then fixated overnight in a 100 mM sodium cacodylate buffer containing 2.5 % (v/v) glutaraldehyde at 4°C. Scanning electron microscopy of the samples was performed at the Biocenter of the Ludwig-Maximilians-University Munich using a Zeiss Auriga cross beam unit (Zeiss, Oberkochen, Germany).

### Amplicon sequencing

Library preparation and sequencing were carried out at the Core Facility Molecular Biology at the Center for Medical Research at the Medical University Graz, Austria. In brief, DNA concentrations were normalized using a SequalPrep™ normalization plate (Invitrogen), and each sample was indexed with a unique barcode sequence (8 cycles index PCR). After pooling of the indexed samples, a gel cut was carried out to purify the products of the index PCR. Sequencing was performed using the Illumina MiSeq device and MS-102-3003 MiSeq^®^ Reagent Kit v3-600cycles (2×251 cycles).

### Sequence data processing and analysis

Demultiplexed, paired reads were processed in R (version 3.2.2) using the R package DADA2 as described in (51). In brief, sequences were quality checked, filtered, and trimmed to a consistent length of ~270 bp (universal primer set) and ~140 bp (archaeal primer set). The trimming and filtering were performed on paired end reads with a maximum of two expected errors per read (maxEE = 2). Passed sequences were de-replicated and subjected to the DADA2 algorithm to identify indel-mutations and substitutions. The DADA2 output table is not based on a clustering step and thus no operational taxonomic units (OTUs) were generated. Each row in the DADA2 output table corresponds to a non-chimeric inferred sample sequence, each with a separate taxonomic classification (ribosomal sequence variants; RSVs) (51). In addition, the merging step occurs after denoising, which increases accuracy. After merging paired end reads and chimera filtering, taxonomy was assigned with the RDP classifier and the SILVA v.123 trainset (52). The visualization was carried out using the online software suite “Calypso” (53). For bar plots data was normalized by total sum normalization (TSS) and for PCoA and Shannon index by TSS combined with square root transformation. Tax4fun was performed based on the Silva-classified OTU table, as described (54).

### RSV Network

G-test for independence and edge weights were calculated on the RSV table using the make_otu_network.py script in QIIME 1.9.1 (55). The network table with calculated statistics was then imported into Cytoscape 3.7.1 (56) and visualized as a bipartite network of sample (hexagons) and RSV nodes (circles) connected by edges. For clustering, a stochastic spring-embedded algorithm based on the calculated edge weights was used. Size, transparency and labels were correlated with RSV abundances, border line intensity refers to RSV persistence over multiple sampling sessions and edge transparency was correlated to calculated edge weights.

### Shotgun metagenomics

Shotgun libraries for Illumina MiSeq sequencing were prepared with the NEBNext^®^ Ultra II DNA Library Prep Kit for Illumina^®^ in combination with the Index Primer Set 1 (NEB, Frankfurt, Germany) according to manufacturer’s instructions and as described in (57). Briefly, 500 ng of dsDNA were randomly fragmented by ultrasonication in a microTUBE on a M220 Focused-ultrasonicator™ (Covaris, USA) in a total volume of 130 μl 1xTE for 80 seconds with 200 cycles per burst (140 peak incident power, 10% duty factor). After shearing, 200 ng of sheared DNA were used for the end repair and adapter ligation reactions in the NEBNext^®^ Ultra II DNA Library Prep Kit for Illumina^®^ according to manufacturer’s instructions. Size selection and purification were performed according to the instructions for 300 to 400bp insert size. Subsequent PCR amplification was performed with 4 cycles and libraries were eluted after successful amplification and purification in 33 μl 1xTE buffer pH 8.0. For quality control libraries were analyzed with a DNA High Sensitivity Kit on a 2100 Bioanalyzer system (Agilent Technologies, USA) and again quantified on a Quantus™ Fluorometer (Promega, Germany). An equimolar pool was sequenced on an Illumina MiSeq desktop sequencer (Illumina, CA, USA). Libraries were diluted to 8 pM and run with 5% PhiX and v3 600 cycles chemistry according to manufacturer’s instructions. Raw fastq data files were uploaded to the metagenomics analysis server (MG-RAST) (58) and processed with default parameters. Annotations of taxonomy (RefSeq) and functions (Subsystems) were then imported to QIIME 2 (2018.11) (59) or Calypso (53) to calculate core features, alpha and beta diversity metrics, statistics and additional visualizations of the datasets.

### Controls

Cultivation, extraction, PCR, and sequencing controls were processed and analysed in parallel to biological samples. An unused wipe not taken out of its bag on the ISS was extracted for every sampling session, cut into pieces, placed on the different media and DNA was also extracted from the solutions obtained with the negative controls. All cultivation controls were negative (no growth of colonies). Wipe solutions of the negative controls used for DNA extraction, PCR, and sequencing revealed a low number of ribosomal sequence variants. These RSVs were removed from according datasets, if present in the samples.

### Data availability

Data are available on request.

## Results

In-flight sampling on board the ISS was performed during increment 51 and 52 (April to June 2017) under the ESA operation named EXTREMOPHILES. All samples (n=24, plus controls) were taken by US astronaut Jack D. Fischer. Three sampling sessions were performed, with session A on 01.05.2017, session B on 12.07.2017 and session C on 29.06.2017. With a span of 72 days between session A and B they were conceptualized for time-course sampling (same sampling locations) to get an idea on the microbiome fluctuation. For comparative purposes, an ISS-relevant cleanroom and a therein housed cargo-spacecraft were sampled, namely cleanroom S5C at the Centre Spatial Guayanais near Kourou in French Guiana with ATV spacecraft.

### ISS microbiome is dominated by human-associated microorganisms and contains also archaeal signatures

The microbial community composition was assessed by amplicon sequence analysis of wipe samples obtained from sessions A-C and the Kourou cleanroom. After complete removal of RSVs from negative controls, more than 3.500 RSVs remained from all ISS and cleanroom samples for the analysis. Both, archaeal and bacterial signatures were retrieved from ISS samples with the universal approach (Fig 1). The signatures belonged to 377 genera, with *Streptococcus, Corynebacterium, Lactobacillus, Acinetobacter, Staphylococcus* predominating (Fig. 1). Firmicutes, Proteobacteria, Actinobacteria and Bacteroidetes were found to be the predominant bacterial phyla (all samples), whereas archaeal signatures (Woesearchaeota, Thaumarchaeota, Euryarchaeota) were frequently detected in air (Cupola air 14.1% and Columbus air session B 3.6% of all sequences) and on various surfaces (Fig. 1). By the archaea-targeting approach, mostly gut associated *Methanobrevibacter* sequences (surface Cupola [sample C2], waste and hygiene compartment [C5]), Woesearchaeota (ATU [A8, B8]; hand grips Columbus [B4]), and unclassified Archaea (ATU [A8], dining table [C7]) were detected. To a lesser extent we also found signatures from Halobacteria (SSC Laptop Columbus [A2]) and Thaumarchaeota (dining table [C7]).

**Fig. 1:**
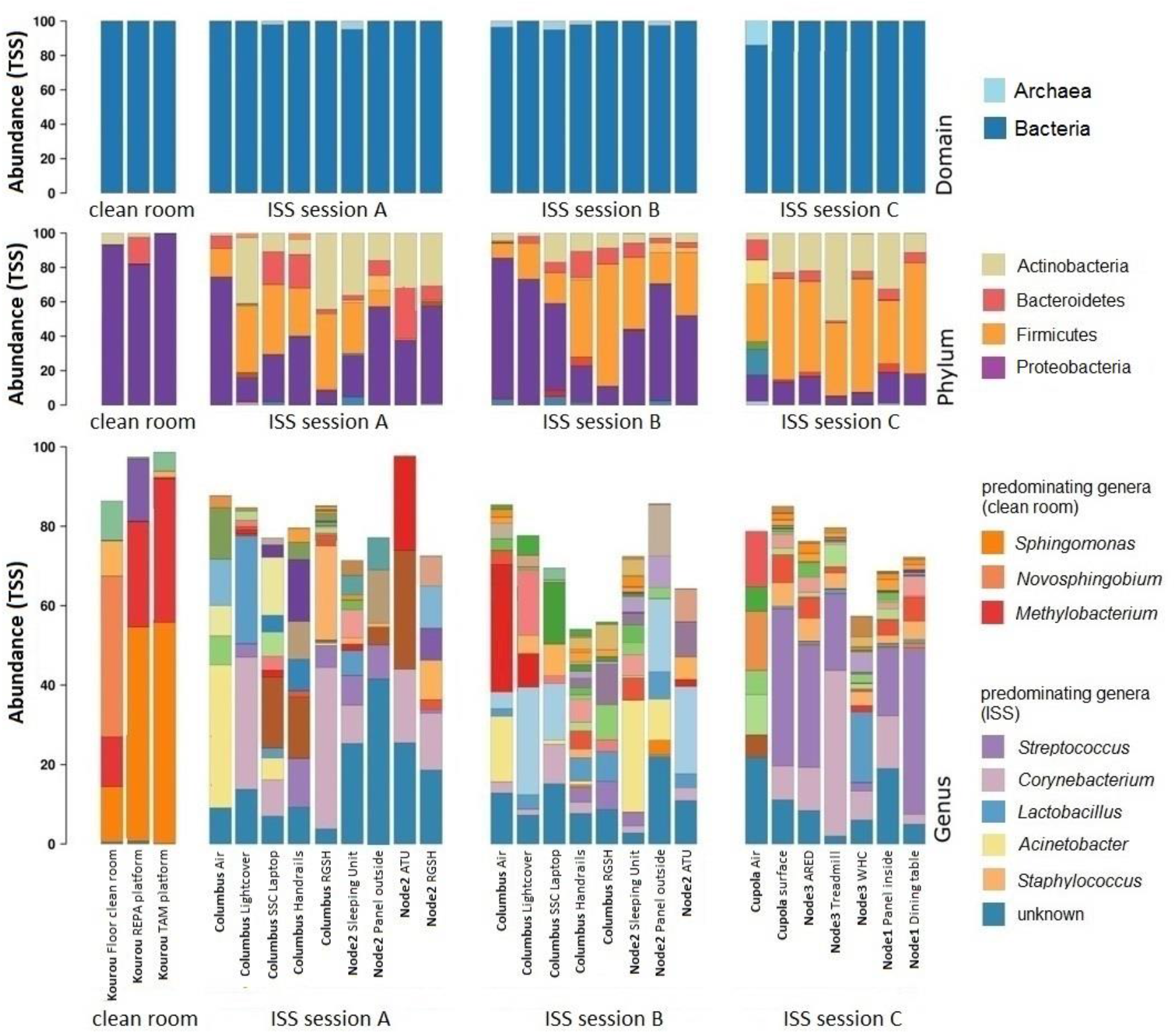
Microbiome composition in cleanrooms (left column) and the ISS sampled during session A, B and C.

The cleanroom samples, which were analyzed for comparison, showed a different microbial signature profile, with a predominance of alpha-proteobacterial genera (*Sphingomonas, Novosphingobium, Methylobacterium*).

### Microbiome dynamics (comparison of session A and B)

To retrieve insights into the microbiome dynamics over time, session A and B were conceptualized with a time-lapse of 72 days between the two samplings. During the complete sampling timeframe, no crew exchange took place, but two cargo deliveries docked (SpaceX and Soyuz). Notably, the microbial diversity (RSVs) were found to be increased significantly in samples of session B (p=0.034, ANOVA; Inverse Simpson’s, rarefied to a depth of 649 RSVs; Fig. 2a), however, the evenness of samples did not change significantly (p=0.68, ANOVA). Anosim analysis indicated a significantly different composition of the samples taken in session A and B (p=0.001). LEfSe analysis (targeting the 300 most abundant genera) identified a substantial increase of signatures belonging to typically gastrointestinal tract-associated genera *Escherichia/Shigella* (p=0.017, ANOVA), *Lachnoclostridium, Ruminococcus_2* (p=0.046) and *Pseudobutyrivibrio* towards session B, whereas members of *Clocibacterium* (p=0.027) and unclassified Corynebacteriaceae (p=0.02) were significantly reduced.

**Fig. 2:**
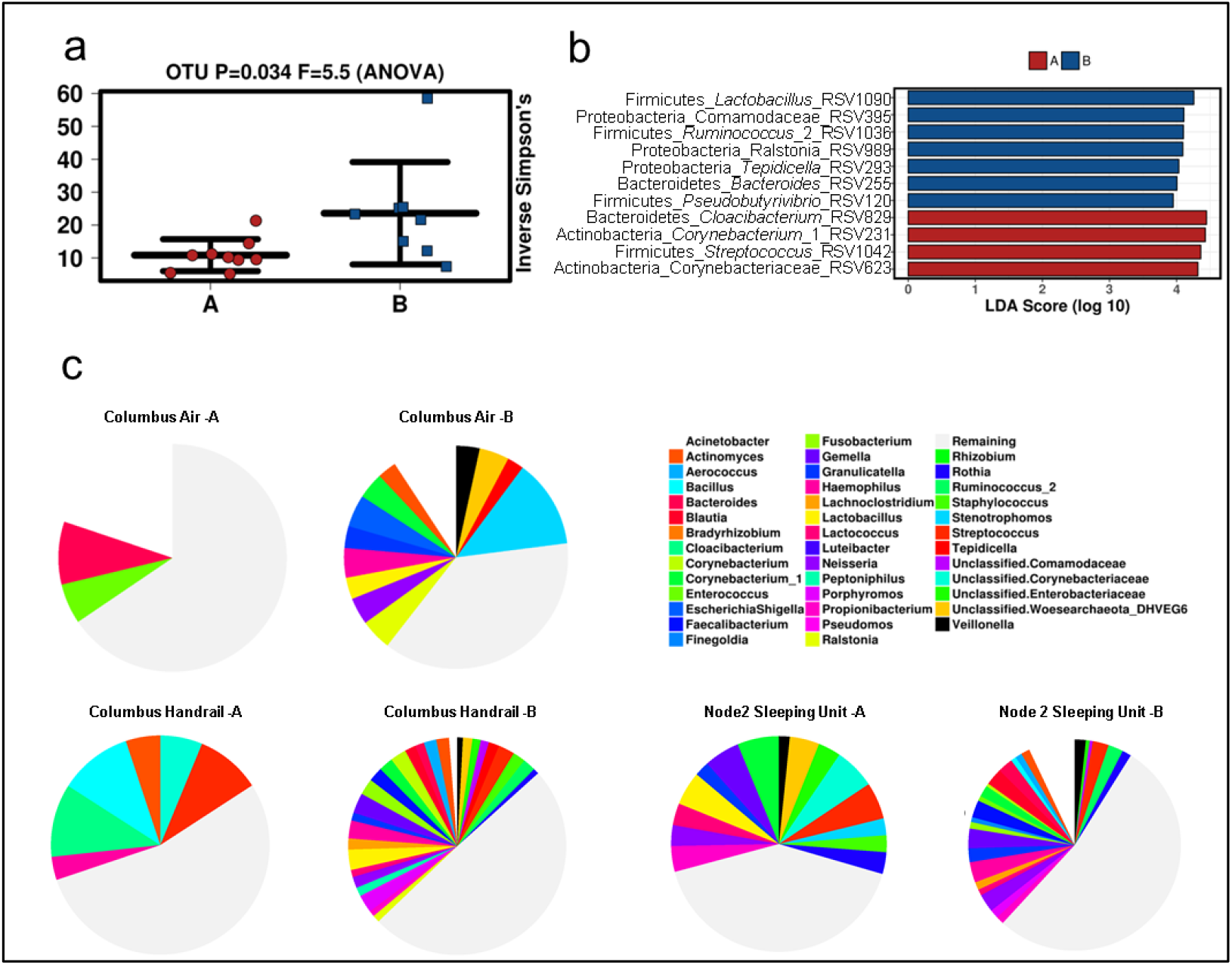
Microbiome diversity and dynamics in samples from sampling session A and B (same locations were sampled, universal approach). a) Inverse Simpson’s index, indicating a significantly different microbiome diversity in samples from session A and B. b) LEfSe analysis (300 most abundant taxa), comparing session A and B. c) Pie charts created including the top 40 most abundant microbial taxa for selected samples. “Columbus Air – A” refers to sample taken from Columbus module: air, in session A.

Significant changes on RSV level are given in Fig. 2b, with an overall notable increase of a certain RSV of *Ralstonia* in samples of session B. Pie charts were created from single locations within the ISS to visualize the changes in microbiome composition (Fig. 2c). It shall be noted, that signatures of unclassified Woesearchaeota (DHVEG6) were found amongst the 40 most abundant microbial genera (additional details below).

### The ISS harbors a core microbiome of more than 50 microbial genera

Core microbiome analyses, looking at the 100 most abundant RSVs, identified 34 taxa shared amongst all sampling time points (A-C, minimal relative abundance: 10%), whereas 55 taxa were shared on genus level. The most abundant, shared RSVs belonged to the microbial genera *Haemophilus, Gemella, Streptococcus, Corynebacterium, Staphylococcus, Lactococcus, Neisseria* and *Finegoldia*. 31 taxa were shared amongst all modules. To obtain a better overview on the biogeography of the ISS microbiome, a network was calculated (Fig. 3).

**Fig. 3:**
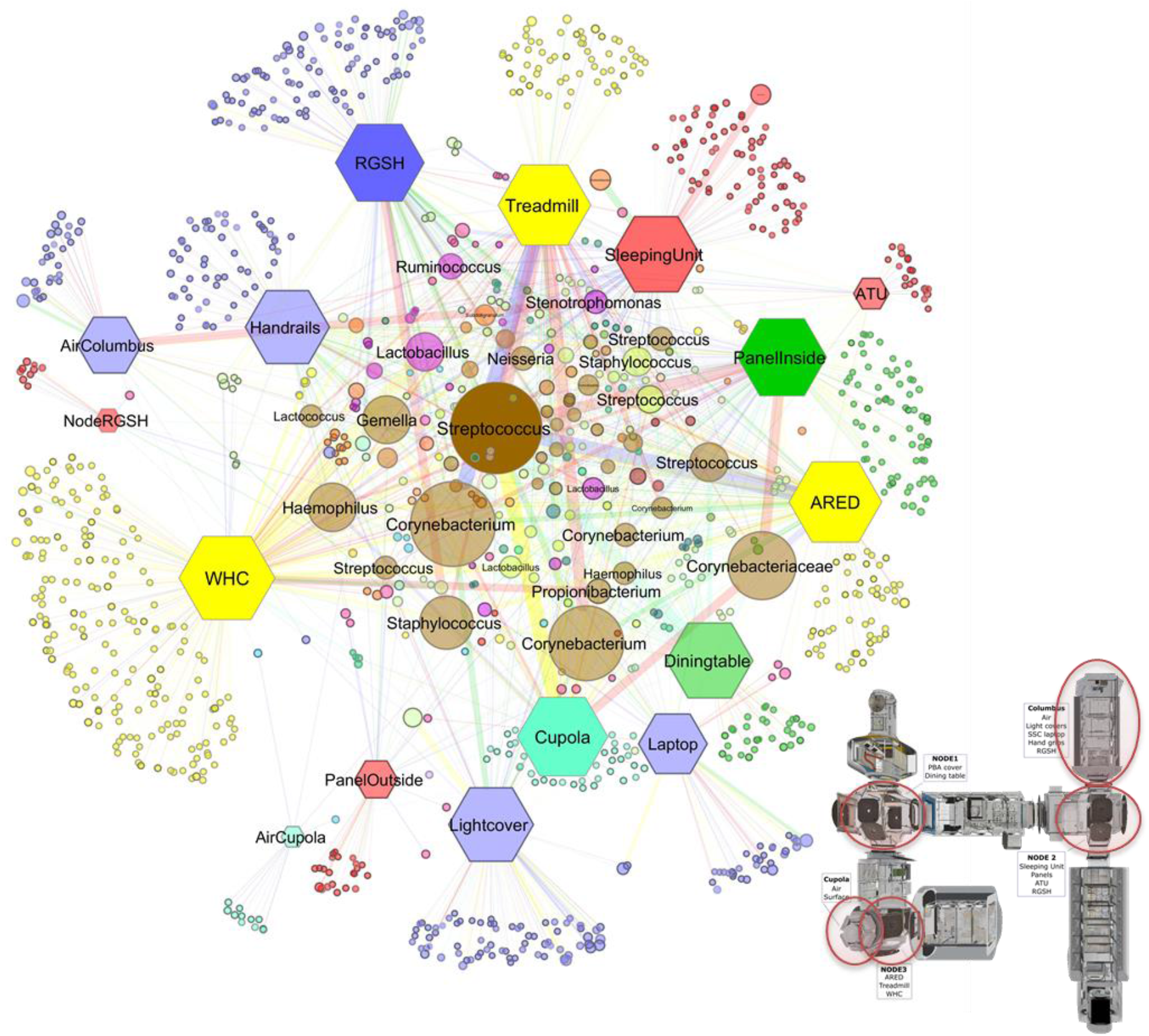
Network analysis and model of the ISS with sampling sites (red circles). Samples are shown as hexagons, RSVs are shown as circles. Color of sample nodes refer to respective ISS modules. RSV nodes show mixtures of these colors if they were shared by several locations and modules of the ISS. The borders of the circles are darker, when found in several sessions. The size, transparency and the size of the node descriptions corresponds to their abundance. The thickness and transparency of the edges (lines) follows the calculated e-weights. Layout: Spring Embedded, with e-weights. Colour code: Yellow (NODE3), red (NODE2), blue (Columbus), aqua (Cupola), green (NODE1). Abbreviations: WHC (Waste and hygiene compartment), ARED (Advanced Resistive Exercise Device), ATU (Audio terminal unit), RGSH (Return Grid Sensor Housing). ISS model source: NASA (https://nasa3d.arc.nasa.gov/detail/iss-internal, accessed on Jan 2019)

The network analysis showed a higher abundance for RSVs which belong to the core ISS microbiome (e.g. *Streptococcus, Corynebacterium, Staphyloccocus, Haemophilus, Gemella* or *Propionibacterium*).

Notably, most location-specific RSVs were observed for WHC (waste and hygiene compartment) and RGSH (return grid sensor housing). This was expected for the WHC area (as hygienic activities shed (internal) human microorganisms into the environment), but surprising for the RGSH. The RGSH is the air inlet part of the air recycling system, expected to accumulate biological burden from the environment, but not to host indigenous microbiology.

Locations with regular crew activity (e.g. treadmill, sleeping unit, handrails) showed higher proportions of RSVs assigned to the human-associated genera *Stenotrophomonas, Ruminococcus* and *Lactobacillus*. When clustering the samples according to their origin, the network also indicated that locations exposed to high human traffic from different modules are more similar to each other than samples of high and low human traffic which were taken within the same ISS module.

### Location shapes microbiome composition

In a next analyses step, we were interested in external parameters influencing the ISS microbiome. Redundancy analysis indicated a significant effect of the time of sampling (sessions; p=0.010), and indicated a potential effect of the location within the ISS (module; p=0.054) on microbiome composition. We further categorized the different samples into: air, personal area (sleeping unit), shared areas which are highly frequented (e.g. communication items, handrails), and shared areas which are less often touched (e.g. lightcover, RGSH, etc.). An NMDS plot performed on RSV level indicated a different composition of the microbiome according to these categories, whereas shared surfaces showed an overlap no matter whether they were frequently or less frequently touched. However, this grouping was not confirmed by statistical analyses (p=0.364, Anosim based on Bray-Curtis distance metric). The highest diversity of microbial signatures was detected in personal areas of the ISS, without being significantly different.

**Fig. 4:**
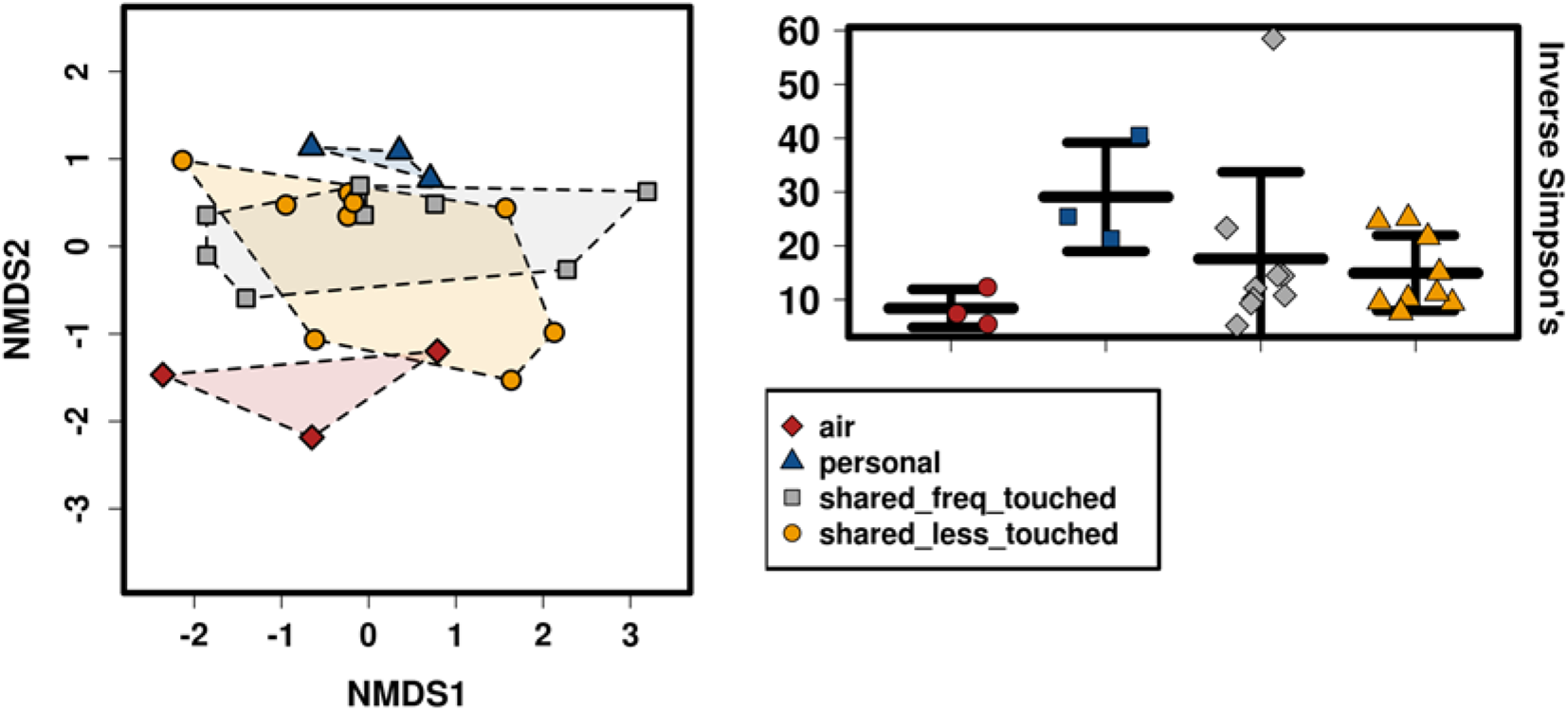
Microbiome composition according to sample categories. Left: NMDS plot (stress 0.129) indicating a separate grouping of personal microbiomes; right: the highest diversity of microbial signatures was observed in microbiome samples from personal areas. The lowest diversity was detected in air samples (p=0.19, Anova; sample depth rarefied to 649 reads).

ANOVA plot analysis indicated e.g. the increased presence of human-associated *Streptococcus* RSVs in samples from dining table and workout area, *Neisseria* species (human mucosa-associated microorganisms) were particularly detected in the sleeping unit and work out area. *Lactococcus* signatures were particularly found in samples from the dining table (potentially food-associated), the sleeping unit and the workout area, whereas RSVs from *Actinomyces, Enterococcus, Lautropia and Brevibacterium* were significantly enriched on handrails, in air, on the dining table, and in the work out area, respectively (all p-values < 0.05, see Fig. 5). The waste and hygiene area showed significantly increased abundances of RSVs belonging to *Lactobacillus, Propionibacterium, Collinsella, Subdoligranulum, Romboutsia* and *Anaerostipes* (Fig. 5). All in all, the samples retrieved from the ISS were largely reflecting the human microbiome, as other sources of the microbial signatures could not be identified (besides potentially food for *Lactococcus*).

**Fig. 5:**
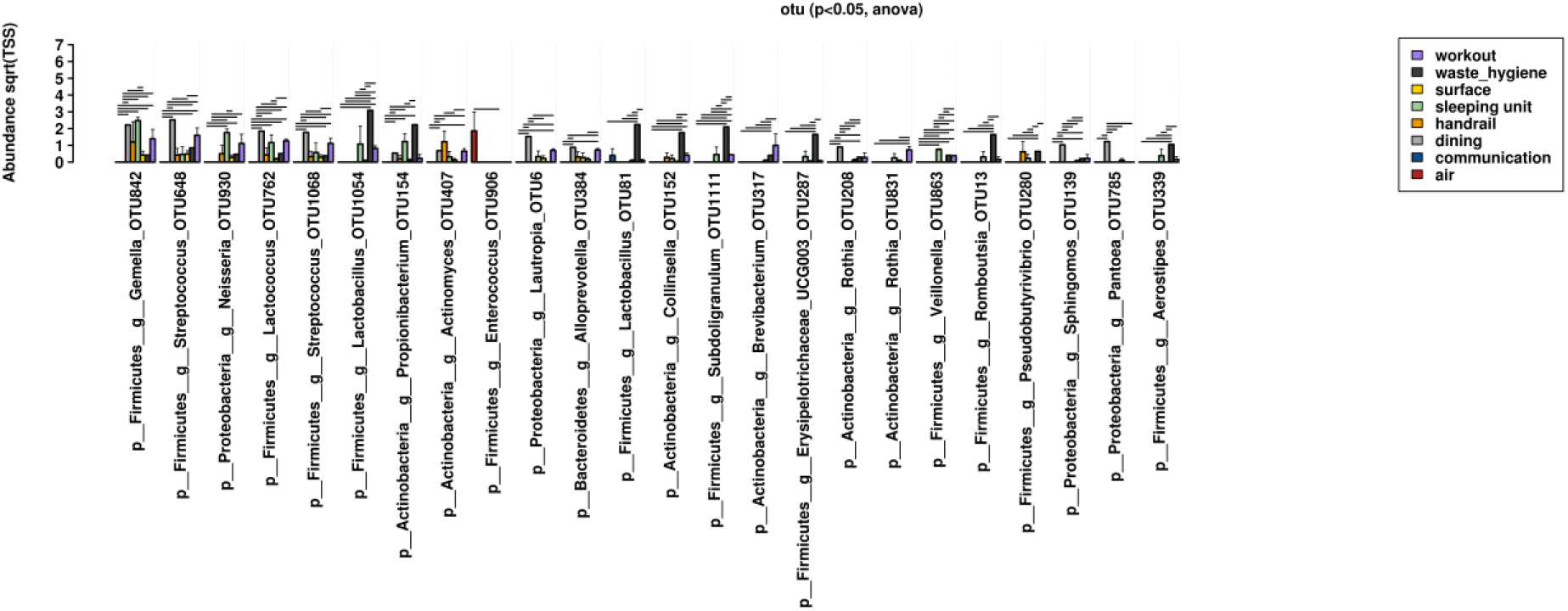
ANOVA plot analysis of different types of locations within the ISS and specifically associated RSVs.

According to a hierarchical cluster analysis based on Pearson’s correlation across session A and B (Fig. 6), a positive correlation of certain microbial phyla with sampled locations was found, being in agreement with findings from the cultivation assays (e.g. *Deinococcus* sp. was isolated from Node2_Panel_Outside). Columbus handrails were found to be correlated with e.g. *Saccharibacteria* and *Bacteroidetes*. A particular pattern was detected for the archaeal signatures retrieved, which were found to be indicative for the sleeping unit (Thaumarchaeota), the handrails (Euryarchaeota) and the samples from the Columbus_SSC_Laptop (Woesearchaeota).

**Fig. 6:**
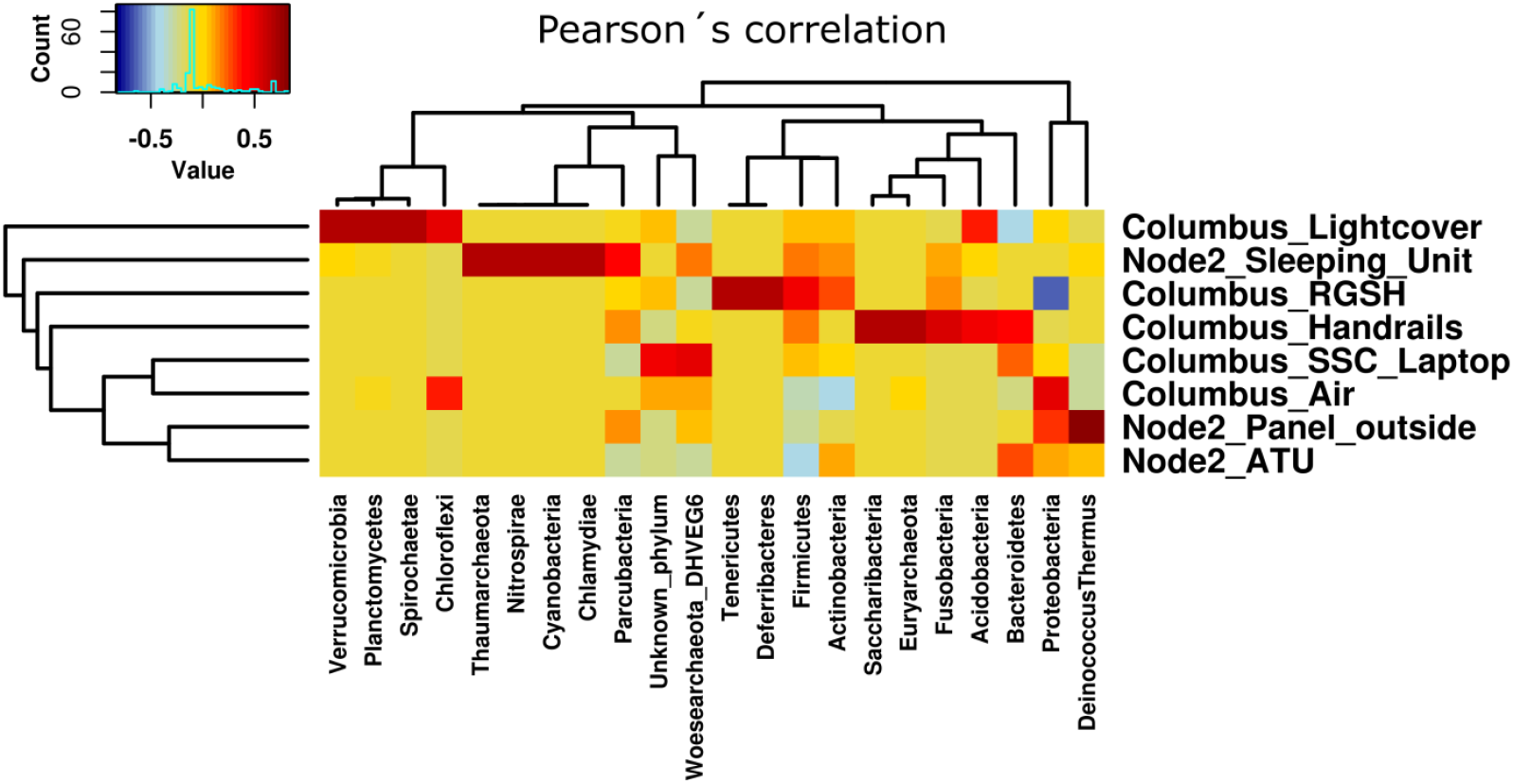
Hierarchical cluster analysis (Pearson’s correlation; universal microbiome data set). Certain microbial phyla (in particular archaeal phyla) were found to correlate with specific sampling sites.

### Microbial functions inferred from 16S rRNA gene information (Tax4fun) and metagenomic analyses

As metagenomics could only be performed on pooled subset of samples (due to low biomass restrictions and sampling set-up, see below), Tax4fun analysis was initially used to predict potential microbial metabolic capabilities, their location-specificity and potential shifts over time.

LEfSe analysis (Fig. 7c) revealed a location specific predicted capacity of the microbial community, with e.g. increased predicted functions in KEGG pathway “Base_excision_repair” in samples from the cupola module, a probably indicator for elevated radiation levels.

**Fig. 7:**
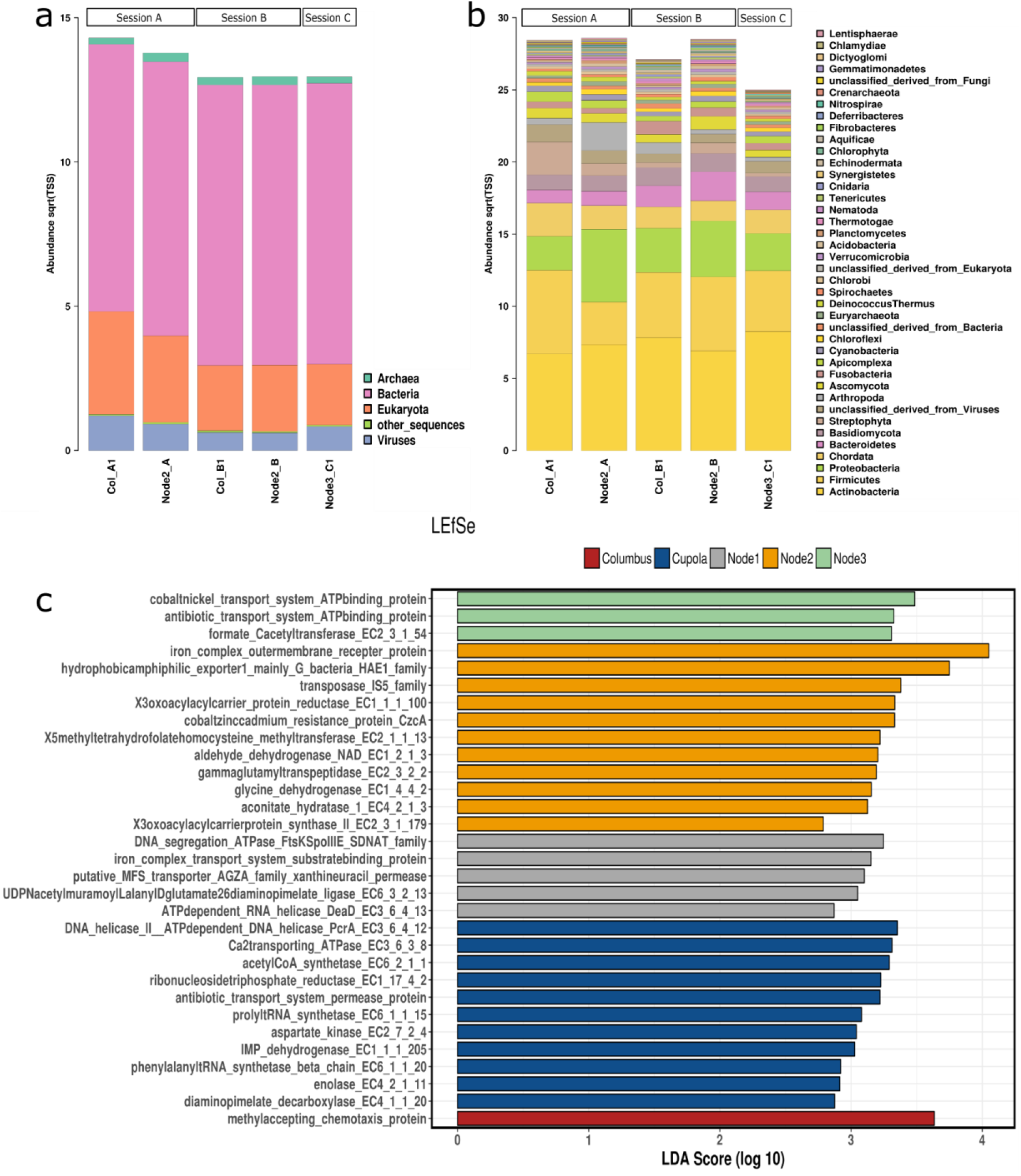
Metagenome-based taxonomic information at kingdom (a) and phylum level (b). LEfSe analysis of predicted functions (Tax4fun), according to module (200 most abundant genetic features; c).

On gene level, different functions were predicted, indicative of a respective module. Node 3 (ARED, treadmill and WHC) revealed predicted signatures of cobalt/nickel and antibiotic transport system ATP binding proteins. Notably, a specific increase of a cobalt/zinc/cadmium resistance protein was predicted for Node 2 (sleeping area and panel samples), an iron complex transport system substrate binding protein for Node 1 (e.g. dining area), and an antibiotic transport system permease protein for the cupola area. This indicated a potential microbial competition and a relevance of transition metal components (e.g. from ISS materials) for the microbial community in these environments.

For metagenomics, samples were pooled as follows: All samples from Columbus module session A (COLA), all samples from Columbus module session B (COLB), all samples Node 1 session C (N1C), all samples Node 2 session A (N2A), all samples Node 2 session B (N2B), all Node 3 session C (N3C).

The taxonomic composition as retrieved from metagenomics was found to be somewhat different to the amplicon-based analysis. In particular the predominance of *Propionibacterium* reads was striking, as its signatures were not well reflected in the amplicon approach. However, *Staphylococcus, Corynebacterium, Streptococcus* could be confirmed as omnipresent on all sampled surfaces of the ISS. However, fungal (e.g. *Malassezia*), viral (e.g. *Microvirus)* and archaeal sequences (e.g. *Methanobrevibacter*) also belong to the core taxa of the ISS (Fig. 7a,b). Core microbial taxa and functions showed a stable distribution over different fractions of samples. Hence, 57% of all taxa and 34% of all functions were shared in all samples (even higher proportions were visible when samples were grouped per module 68% and 51% or per sampling session 73% and 58% respectively).

Sample N2B showed the highest Shannon diversity on genus (H’ ~ 4) and functional level (H’ ~ 10). Similarity estimates based on Bray-Curtis distances revealed that microbes and their functions from the Columbus module showed the biggest difference to samples from Node 1, 2 and 3. In addition the Columbus module also experienced the biggest shift of its microbiome along PCoA Axis 1 (taxa: ~60%, functions ~40%) between the first and last sampling session.

Regarding functions, genes assigned to arginine biosynthesis (amino acid metabolism), degradation of L-Ornithine (amino acid metabolism), copper-translocating P-type ATPases (virulence, disease and defense), and the phage major capsid protein (phages, prophages, transposable elements) were ubiquitously distributed, whereas functions assigned to dormancy and sporulation, photosynthesis, motility and chemotaxis as well as aromatic compounds metabolism showed location-dependent variations. Functions involved in iron acquisition and metabolism (ferrous iron transport protein B: 0.2% in functional core), potassium metabolism, nickel ABC transporters and others were highly abundant, indicating a potential surface-interaction with ISS materials.

### ISS cultivable microbial community reveals extremo-tolerant traits

In the course of this study, hundreds of colonies/cultures retrieved on/in various media were processed, resulting in 76 unique bacterial isolates. Along with the bacteria, also fungi were isolated, but are not further analyzed herein. These included *Aspergillus* species *(A. sydowii, A. unguis), Chaetomium globosum, Penicillium* species (*P. aurantiogriseum, P. brevicompactum, P. chrysogenum, P. crustosum, P. expansum*), *Rhizopus stolonifera* and *Rhodotorula mucilaginosa*. All fungal isolates obtained in this study were assigned to biosafety risk group S1. *P. brevicompactum, P. chrysogenum, P. crustosum, R. stolonifera* and *R. mucilaginosa* may cause allergenic reactions and *R. mucilaginosa* can also act as opportunistic pathogen. Archaea could not be grown from any sampling site.

Most of the bacterial isolates showed a distinct pattern in origin distribution and special growth/enrichment characteristics, which is shown in Fig. 8.

**Fig. 8:**
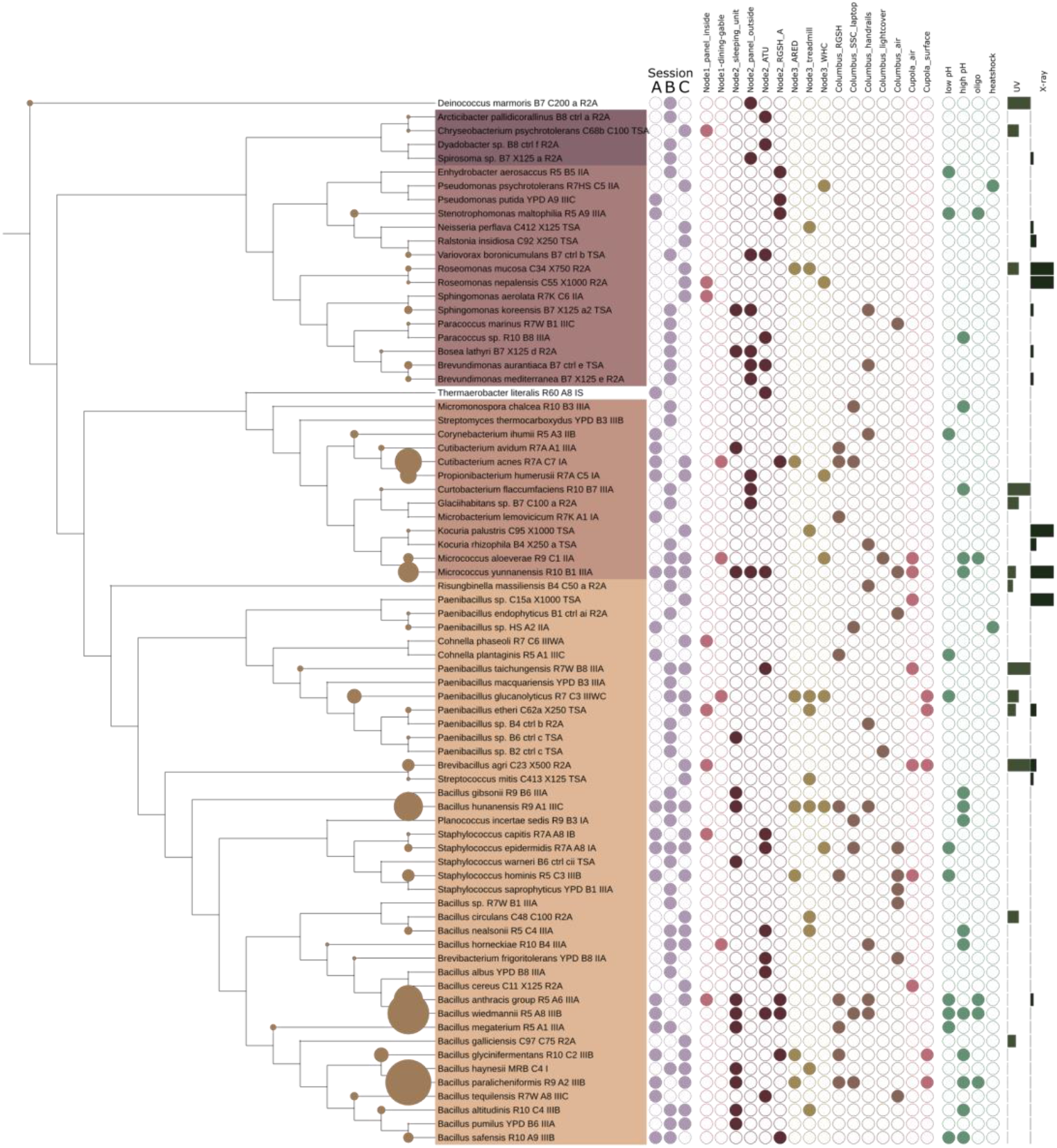
Tree of all isolates retrieved from sessions A-C. Circles on branches reflect the number of isolates obtained from the respective species. First dataset reflects the session number (A-C), the locations where the isolates were retrieved from (Node 1, 2, 3, Columbus and Cupola in different colors), the enrichment conditions (low, high pH, oligotrophic conditions and heat-shock resistant) and the radiation dose (UVC, X-Ray; bars on the right) which were used to pre-treat the samples.

It shall be noted, that a number of isolates was obtained under stringent cultivation or pre-treatment conditions. This included i) UV- and X-ray resistant microorganisms, such as *Deinococcus marmoris, Curtobacterium flaccumfaciens, Brevibacillus agri* (UV_254 nm_: 200 J/m^2^), *Roseomonas* species, *Kocuria palustris, Micrococcus yunnanensis, Paenibacillus* sp. (X-ray: 1000 Gy), ii) microorganisms growing particularly at high or low pH, or iii) heat-shock survivors (*Pseudomonas psychrotolerans, Paenibacillus* sp.) (Fig. 7). Isolates retrieved under non-mesophile conditions were, for example, *Thermaerobacter literalis* (a true thermophile isolated at 65°C from the ATU in Node2, does not grow below 50°C), *Sphingomonas aerolata* and *Microbacterium lemovicicum* (exhibiting extraordinary cryotolerance, isolated only at 4°C, maximal growth temperatures were 51°C and 32°C, respectively).

### Physiological characteristics and resistance potential of ISS microbes do not differ from ground controls

In the next step, we analyzed physiological characteristics of ISS microbial isolates. In particular, we were interested whether they withstand physical and chemical stressors better than same or closely related microbial species from ground controls.

For these tests, we selected a subset of microbial isolates from the ISS, spanning 11 microbial genera of interest (listed in Fig. 9). This list included typical confined-indoor bacteria, like *Bacillus, Micrococcus* and *Staphylococcus*, but also microorganisms of special interest (associated to spacecraft assembly, extraordinary hardy, extremophile) were included (e.g. *Microbacterium, Cupriavidus*, or *Ralstonia*). For comparative reasons, we included also eight microbial isolates from ground controls (cleanrooms, Concordia station) or culture collections. Overall, the final list comprised 29 different microbial strains.

**Fig. 9:**
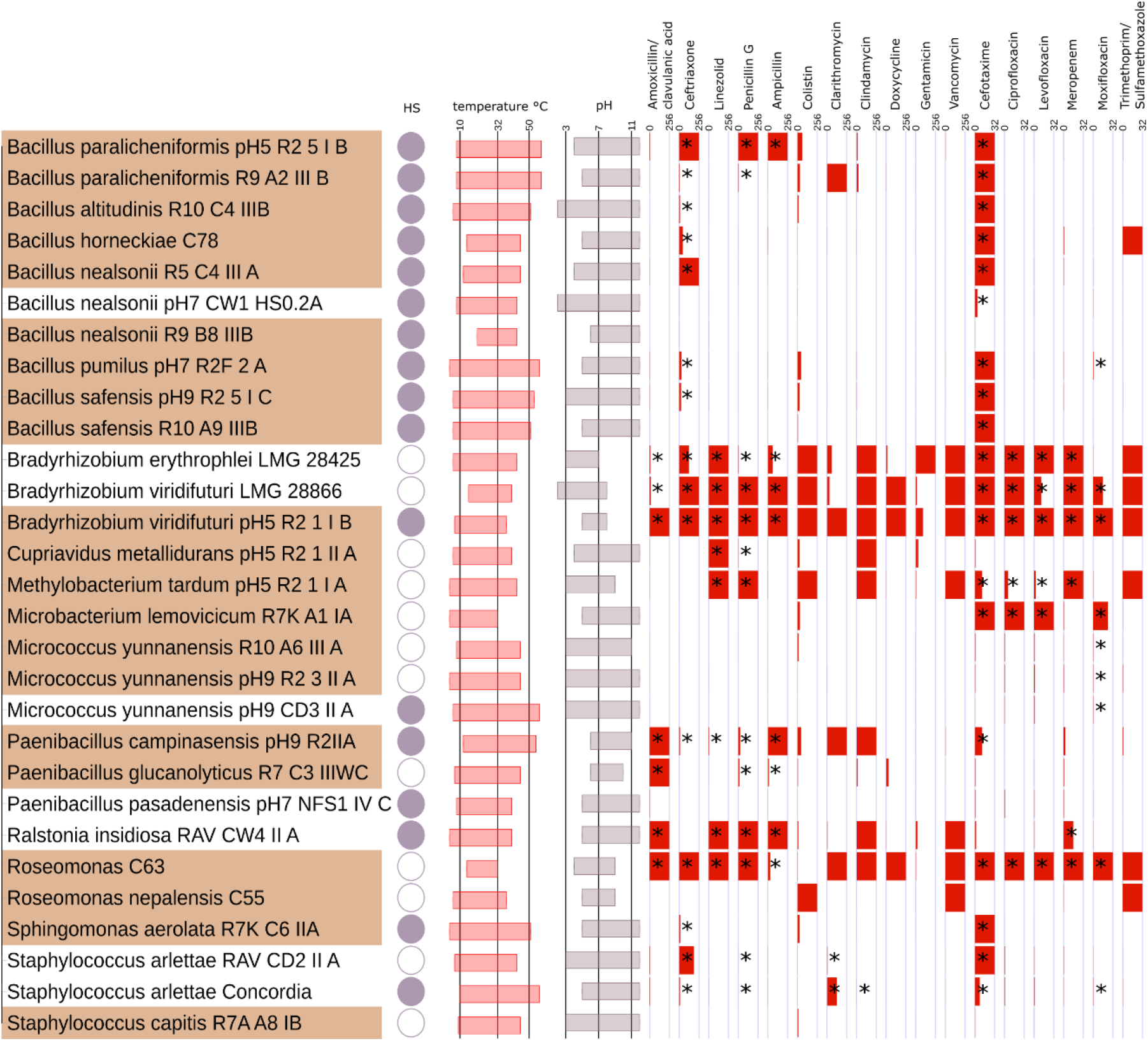
Selected microorganisms from ISS (highlighted in brownish color) and closely related isolates from ground controls (cleanrooms, culture collections, Concordia station; white). Datasets reflect resistances to heat shock (filled circle: heat-shock resistance), temperature range of growth, pH range of growth and resistance towards antibiotics. Antibiotics were applied in maximal concentrations of 256 and 32μg/ml, as indicated. Bars reflect the minimal inhibitory concentrations as determined experimentally. A full red bar indicates resistance against the maximal concentration tested. Asterisks indicate when an isolate was deemed resistant against a certain antibiotic according to the EUCAST guidelines for PK/PD (non-species related) and *Staphylococcus* spp. breakpoints. Partial information on some of the isolates is already available in (14)

All these strains were tested with respect to heat-shock resistance in the stationary phase (*Bacillus* cultures contained spores), upper and lower temperature limit (growth), upper and lower pH limit (growth), and resistance towards a variety of antibiotics (Fig. 9).

Antimicrobial susceptibility testing for 17 clinically relevant antibiotics was performed. Antibiotic resistance/susceptibility was found to be in some cases strain-specific but mostly species/genus-specific, independent from their isolation source (ISS or ground control). In particular the tested *Bradyrhizobium* species showed a vast resistance against numerous antibiotics, as did one *Roseomonas* strain. The antibiotic resistance pattern was judged following the EUCAST guidelines for PK/PD (non-species related) or, for *Staphylococcus* isolates, *Staphylococcus* spp. breakpoints (14) were used. It has to be stressed that all tested isolates were non-pathogenic and that these results shall not be used for clinical risk assessment of any kind. The ISS strains were not found to be significantly more resistant (number of antibiotics or concentration) than their counterparts from ground controls.

Notably, the strains showed a growth temperature span of 18 to 52 degrees (14-32°C, *Roseomonas* C63; *Bacillus pumilus*, 4-56°C). The minimal and maximal growth temperatures, or the temperature span, were not significantly different in ISS isolates compared to control microorganisms (Mann-Whitney U Test). For growth at different pH values, the isolates revealed a pH span of 4 to 10 pH values *(Bradyrhizobium erythroplei* LMG28425, pH3-7; *Bacillus altitudinis* R10_C4_IIIB, pH 2-12). As seen for the temperature, no significant difference in pH preference of ISS strains versus ground control strains was observed (Mann-Whitney U Test).

### Genomic properties of ISS isolates showed no differences to non-ISS neighbors, but revealed partial discrepancy of genomic and experimentally observed antibiotics resistance patterns

In order to understand the specific genomic characteristics of ISS microorganisms, we selected six different species for genome sequencing and reconstruction, namely: *Bacillus pumilus* strain pH7_R2F_2_A, *Bacillus safensis* strain pH9_R2_5_I_C. *Bradyrhizobium viridifuturi* strain pH5_R2_1_I_B, *Cupriavidus metallidurans* strain pH5_R2_1_II_A, *Methylobacterium tardum* strain pH5_R2_1_I_A and *Paenibacillus campinasensis* strain pH9_R2IIA (41) and compared the assemblies to publicly available genomes.

The genome of *<u>Bacillus pumilus</u>* strain pH7_R2F_2_A was retrieved 99.59% complete, with a %GC of 41.6. The overall genome length was 3.7 Mbp. *Bacillus pumilus* SAFR-032 (3.7 Mbp, 41.3 %GC; ENA study ID: PRJNA20391), whose genome was analysed for comparative reasons as well, possessed the same antibiotics resistance capacity. The ISS strain possessed all necessary genes for flagellum assembly and CAS-TypeIIIB (with cmr5_TypeIIIB missing); the latter was not found in *Bacillus pumilus* SAFR-032. Looking at the metabolic profiles, the ISS isolate of *Bacillus pumilus* (comparison to SAFR-032 and ATCC 7061 (3.8 Mbp, 41.7 %GC; ENA study ID: PRJNA29785) possessed the genomic capacity to perform choline and methionine degradation, but no other peculiarities were identified.

The genome of *<u>Bacillus safensis</u>* strain pH9_R2_5_I_C was found to be 99.59% complete, with a %GC of 41.5. The overall genome length was 3.7 Mbp. It possessed all necessary genes for flagellum assembly and CAS-TypeIIIB, as did next neighbour *Bacillus safensis* FO-36b. Looking at the metabolic profiles, the ISS isolate of *Bacillus safensis* (comparison to CFA06 (3.7 Mbp, 41.5 %GC; ENA Study ID: PRJNA246604) and FO-36b (3.7 Mbp, 41.6 %GC; ENA Study ID: PRJNA270528) did not show certain peculiarities.

The genome of *<u>Bradyrhizobium viridifuturi</u>* strain pH5_R2_1_I_B was found to be 99.96% complete, with a %GC of 64.3. The overall genome length was 7.9 Mbp. The genome carried several copies of the efflux pump membrane transporter BepG as well as other multidrug efflux transporters and β-lactamase genes, which largely explained the overall stable antibiotic resistance observed in our experiments. The observed resistances against linezolid and vancomycin could not be directly inferred from the genomic data. These features were also found in *Bradyrhizobium viridifuturi* SEMIA 690 (8.8 Mbp, 64.0 %GC; ENA Study ID: PRJNA290320), the next phylogenetic neighbour. Overall, the genetic features of our ISS isolate were widely similar to known *Bradyrhizobium* species. The only differences found were the potential capability for homospermidine biosynthesis from putrescine and (R)-acetoin biosynthesis through (S)-2-acetolactate.

The genome of *<u>Cupriavidus metallidurans</u>* strain pH5_R2_1_II_A was found to be 99.94% complete, with a GC content of 63.7 %. The overall genome length was 6.9 Mbp. This strain carries three bepE efflux pump membrane transporters, and also a multidrug efflux system protein (acrB). However, the bepE efflux pumps were not detected in the genome of its closest relative *C. metallidurans* CH34. The genome showed full potential for type IV pili and flagella formation and numerous secretion systems, but this was not a unique feature for the ISS strain. With respect to the metabolic profile, *C. <u>metallidurans</u>* strain pH5_R2_1_II_A showed a number of different features when being compared to the next relatives (*C. basilensis* OR16, ENA Study ID: PRJNA79047; *C. metallidurans* CH34, ENA Study ID: PRJNA250; *C. necator* N-1, ENA Study ID: PRJNA67893; *C. taiwanensis* LMG19424, ENA Study ID: PRJNA15733), which included the predicted capacity for 5,6-dimethylbenzimidazole biosynthesis and butanediol degradation/synthesis.

The genome of *<u>Methylobacterium tardum</u>* strain pH5_R2_1_I_A was found to be 100% complete with a GC content of 69.2%, and a total genome length of 6.5 Mbp. Also *M. tardum* pH5_R2_1_I_A carried the efflux pump membrane transporter BepE and the genetic capacity for flagellum formation and several secretion systems. The strain showed a number of differential features when we compared the genomic potential with other members of the genus (M. *extorquens, M. mesophilicum, M. nodulans, M. populi, M. radiotolerans;* 5,6-dimethylbenzimidazole biosynthesis, base-degraded thiamine salvage, cytidylyl molybdenum cofactor biosynthesis, L-dopachrome biosynthesis); however, it shall be noted, that another genome of the species was not available at the time of analysis.

The genome of *Paenibacillus campinasensis* strain pH9_R2IIA could be retrieved with a 99.84% completeness. It showed a GC content of 52.26%, and a genome length of 5.4 Mbp. The genome revealed a potential for lincosamide (Clindamycin), macrolide (clarithromycin), fluorquinolone (moxifloxacin, levofloxacin, ciprofloxacin), and glycopeptide (vancomycin) resistance which could all be verified by the antimicrobial susceptibility tests with the exception of the vancomycin resistance (no PK/PD breakpoint in the EUCAST table). Nevertheless, the observed MIC for vancomycin was 4 μg/ml, which was the highest observed MIC for vancomycin besides the seven isolates which were completely resistant (see Fig. 8). The genome did not show any β-lactam resistances but in spite of this, *Paenibacillus campinasensis* strain pH9_R2IIA was resistant against all β-lactam antibiotics with the exception of meropenem in the antimicrobial susceptibility tests. The strain showed the potential for flagella formation, and the presence of CAS type III. At the time of the analysis there was no other genome of this species publically available, but the metabolic potential was not found to be strikingly different from other genome-sequenced members of the *Paenibacillus* genus.

The antibiotic resistance genes (ARG) detected in the sequenced genomes and the inferred antibiotic resistances are summarized in Fig. 10. These detected ARGs conformed for the most part with the results from the antimicrobial susceptibility tests; however, there were also some discrepancies. For example, both *Bacillus* strains had genes for the transcription-repair coupling factor mfd and the efflux transporter blt which should provide resistance against multiple fluorquinolones, but *Bacillus pumilus* strain pH7_R2F_2_A was only resistant against moxifloxacin and not against ciprofloxacin or levofloxain, while *Bacillus safensis* strain pH9_R2_5_I_C was sensitive against all tested fluorquinolones. *C. metallidurans* was unharmed by the lincosamide clindamycin and the oxazolidinone linezolid and grew at the maximal tested concentrations of these antibiotics, but these resistances could not be inferred from the ARG’s. Moreover, *C. metallidurans* was sensitive to all fluoroquinolones and β-lactam antibiotics besides Penicillin G in spite of possessing several efflux transporter genes from which a resistance against fluorquinolones can be inferred and also the β-lactamase AmpC which is a specialized cephalosporinase and infers resistance against the tested cefotaxime and ceftriaxone. However, it is known that these two cephalosporins, while being sensitive to AmpC, are only weak inducers for actual AmpC expression (60).

**Fig. 10:**
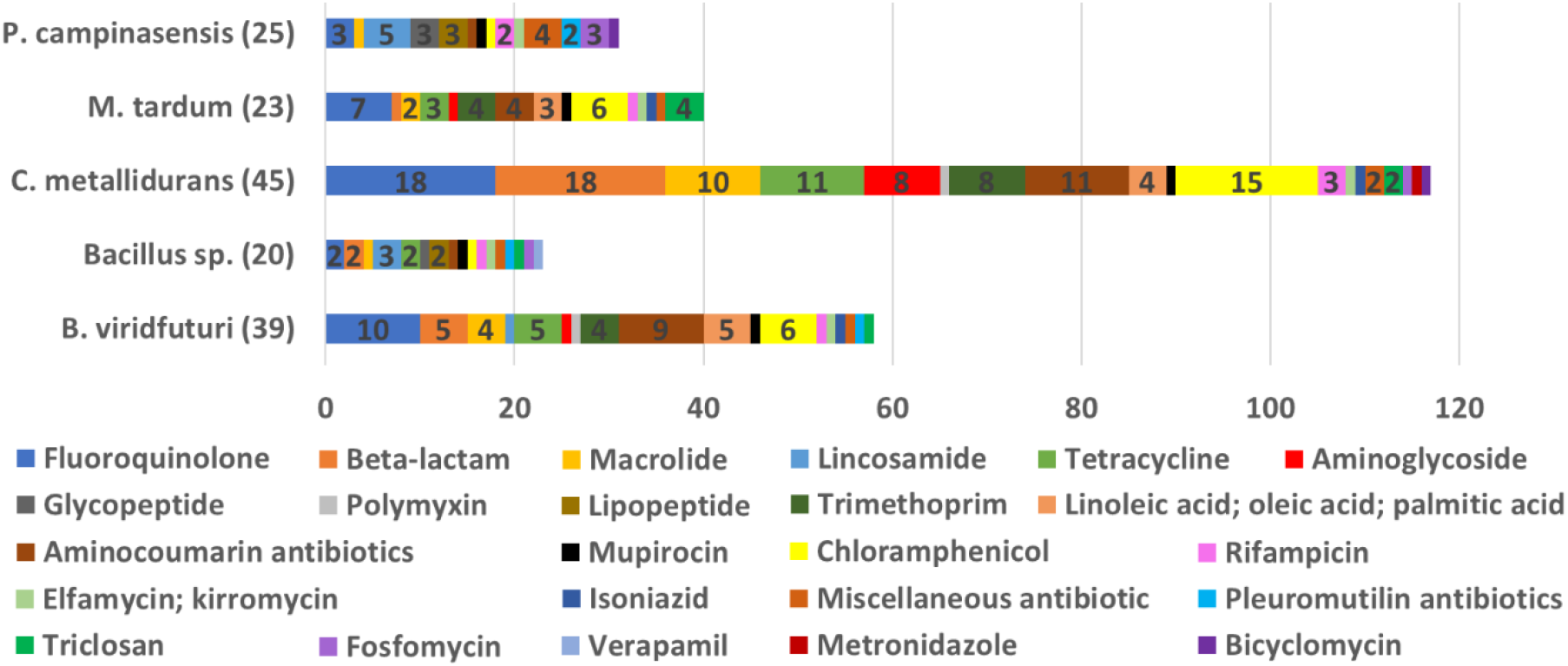
Summary of the antibiotic resistance genes (ARG) detected in the sequenced genomes. The number in () is the total number of detected ARG in the respective genome and the colored columns show the groups of antibiotics against which the detected ARG are known to provide resistance. Numbers within a colored column indicate how many of the detected ARG provide resistance against this type of antibiotic; in case of one ARG was found, no number is given. As some ARG, especially the multidrug efflux pumps which were found in a high number in *C. metallidurans* and *B. viridfuturi*, may be able to provide resistance against multiple kinds of antibiotics, the sum of the numbers within the colored columns may exceed the total number of detected antibiotic resistance genes per organism. The number of ARG as well as of inferred antibiotic resistances was the same in the sequenced *Bacillus* genomes, which is why they are summarized as one *Bacillus sp*. in this graph.

### The ISS microbiome in context: humans and the cleanroom as a contamination source

The microbiome of an ISS cargo-harboring cleanroom, which we analyzed for comparative reasons, was found to be different from the ISS microbiome. The microbial diversity detected in cleanrooms was significantly lower than observed in ISS samples (ANOVA; p=0.012; Shannon diversity index) and clustered separately in multivariate analyses (Fig. 11). The cleanroom microbiome was specifically characterized by a predominant abundance of α-Proteobacteria such as Novosphingobium, *Sphingomonas* and *Methylobacterium* whereas most ISS samples were dominated by Firmicutes and Actinobacteria. This is in accordance with previous findings (13). Based on this observation, we argue that the items delivered from terrestrial cleanrooms to the ISS are most likely not a (relevant) microbiome source.

**Fig. 11:**
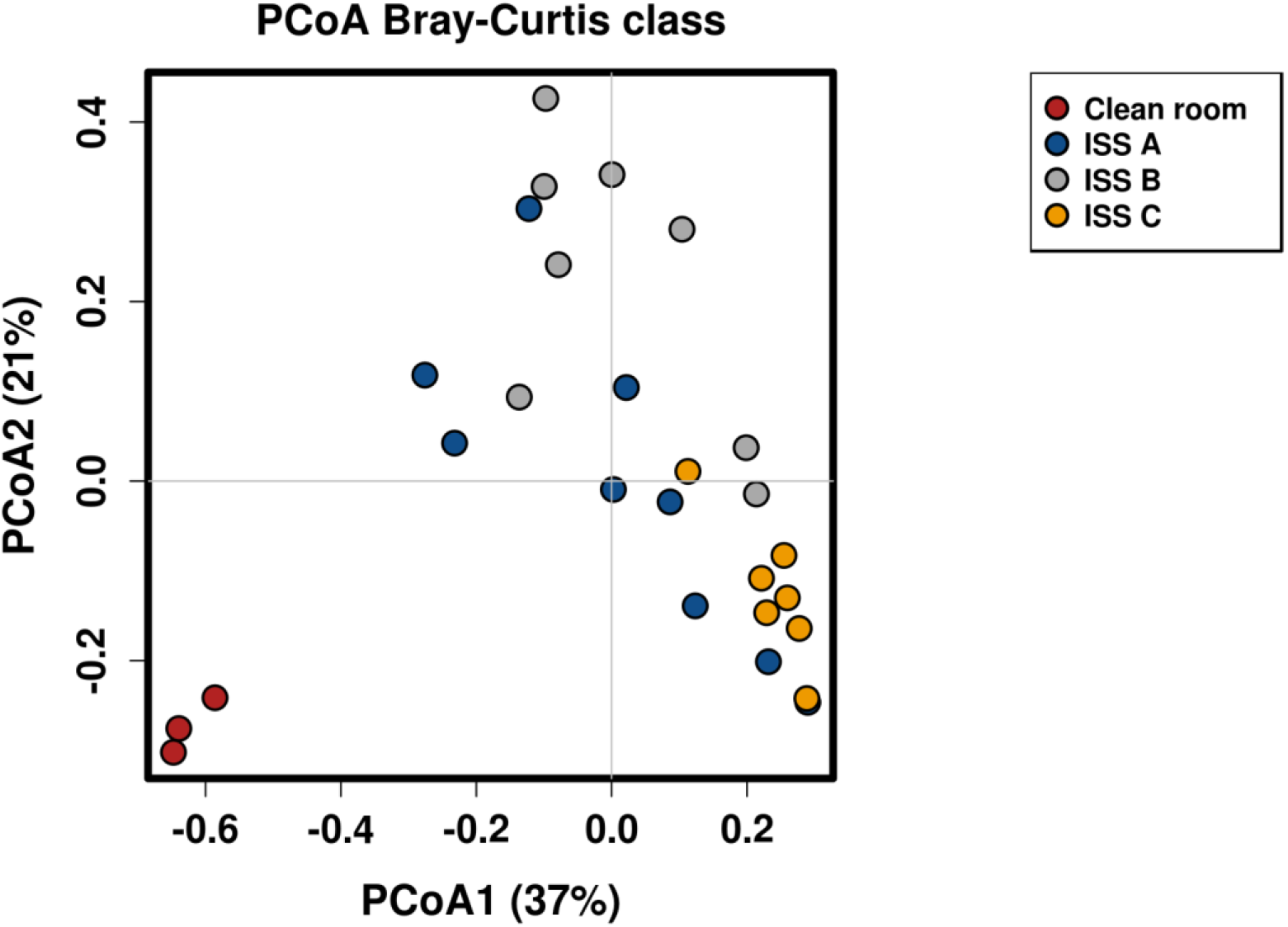
PCoA plot or microbiome compositions of cleanroom and ISS samples

However, a more detailed picture was obtained, when we looked at the cultivable diversity retrieved from ISS and cleanroom samples (Fig. 12), where we found an overlap of several bacterial species, including: *Bacillus cereus, B. aerophilus, B. subtilis, B. nealsonii, Micrococcus aloeverae, M. yunnanensis, Kocuria palustris, Ralstonia insidiosa*. Human-associated microorganisms, such as e.g. *Micrococcus* species, were more likely introduced by humans into both environments than transported via cargo from cleanrooms to the ISS. Nevertheless, this comparison indicates a potential transfer of cleanroom associated microorganisms (such as *Bacillus* species) onto the ISS where they established themselves as a part of the ISS microbial community. Based on the total cultivable diversity they comprise only a minor fraction of the ISS microbial community.

**Fig. 12:**
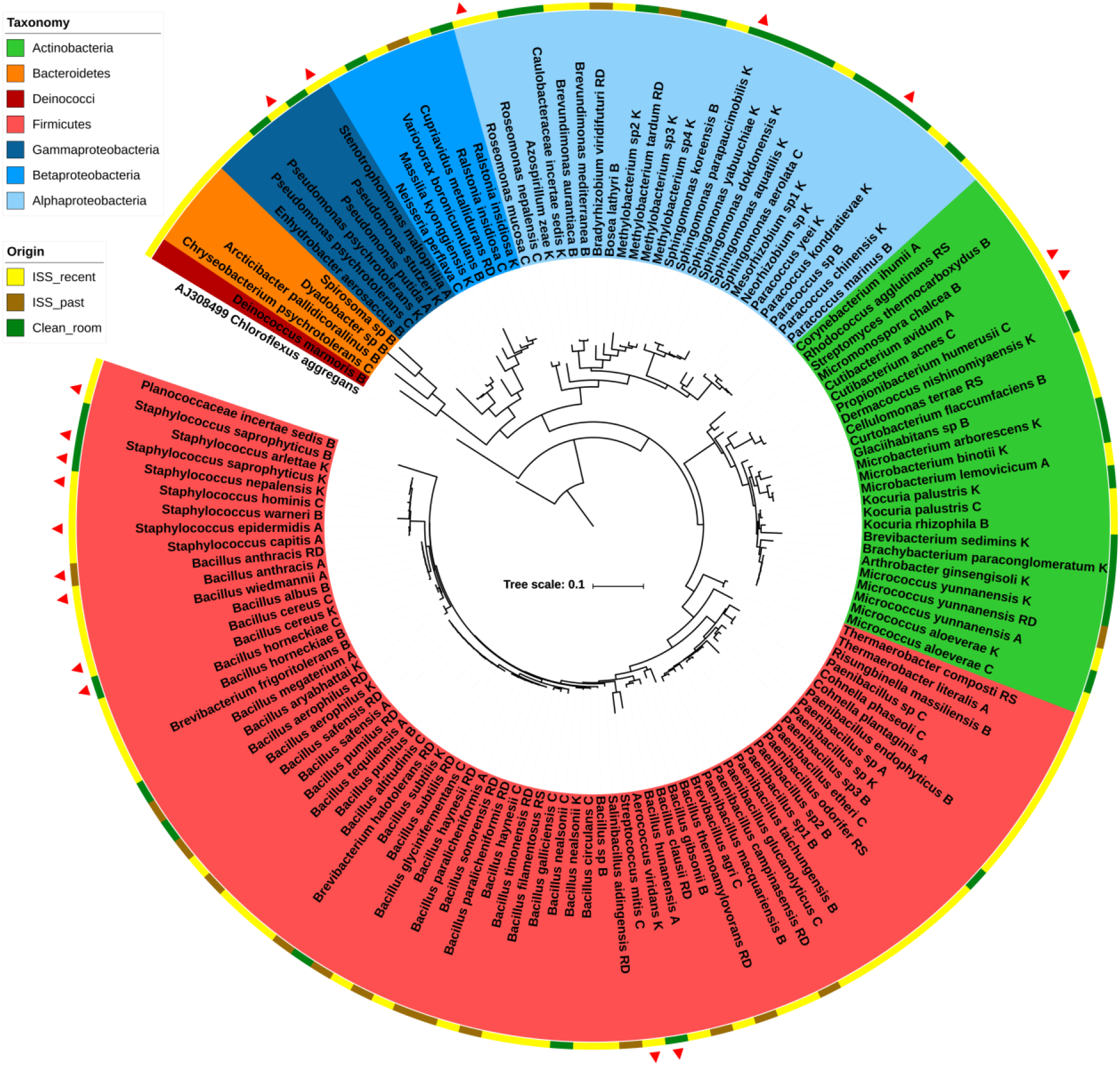
Phylogenetic tree of ISS isolates from this (“ISS_recent”) and our previous study (14) (“ISS_past”), including cleanroom isolates obtained during the Kourou/ ATV sampling campaign (this study; “Clean_room”). Red triangles indicate a biosafety risk classification of S2. Isolates which are not assigned a complete species name had less than 98.5% 16S rRNA gene sequence similarity to their closest described neighbour.

### The ISS is a source of novel microbial species

The application of 21 different cultivation approaches resulted in a high diversity of microbial isolates from the ISS, and 22 of the bacterial genera obtained during this study have not been isolated from ISS samples before, although most of them have been detected by molecular methods (12,13,19) (*Arcticibacter, Bosea, Brevundimonas, Chryseobacterium, Cohnella, Curtobacterium, Cutibacterium, Deinococcus, Dyadobacter, Enhydrobacter, Glaciihabitans, Micromonospora, Neisseria, Paracoccus, Planococcus, Propionibacterium, Risungbinella, Roseomonas, Spirosoma, Stenotrophomonas, Thermaerobacter*, and *Variovorax*). Of our fungal isolates, only *Aspergillus unguis* was not found on the ISS before.

Additionally eleven of the ISS isolates obtained in this study might even qualify to comprise novel, hitherto undescribed bacterial species as their 16S rRNA gene sequence similarity to their respective closest described neighbor was below 98.5%. Six isolates thereof revealed a similarity lower than 97%, including *Paenibacillus, Dyadobacter, Spirosoma* representatives. The isolate with the highest nucleotide difference was isolate R9_B3_IA sampled from the SSC laptop in the Columbus module, as closest neighbour was *Planococcus halocryophilus* (87.36% sequence identity), an extreme psychrophile isolated from arctic permafrost (61).

### Do ISS microorganisms interact with ISS surfaces? Co-incubation experiments with ISS materials

ISS isolates *Cupriavidus metallidurans* strain pH5_R2_1_II_A, *Bacillus paralicheniformis* strain R2A_5R_0.5 (14) and *Cutibacterium avidum* strain R7A_A1_IIIA were aerobically and anaerobically incubated together with untreated aluminium alloy platelets, eloxated aluminium alloy platelets, and pieces of NOMEX^®^ fabric, to investigate if these isolates interact with these ISS surface materials. After incubation (6 weeks; 32°C; liquid R2A medium), the co-incubated materials were analyzed via scanning electron microscopy (Fig. 13). The NOMEX^®^ fabric itself remained intact over time (Fig. 13, I-L, negative control), but served as an excellent attachment surface for *B. paralicheniformis* biofilms and single cells of *C. metallidurans*. The ability of *C. metallidurans* to attach to surfaces was found genetically encoded by pili formation capacity (see above).

**Fig. 13:**
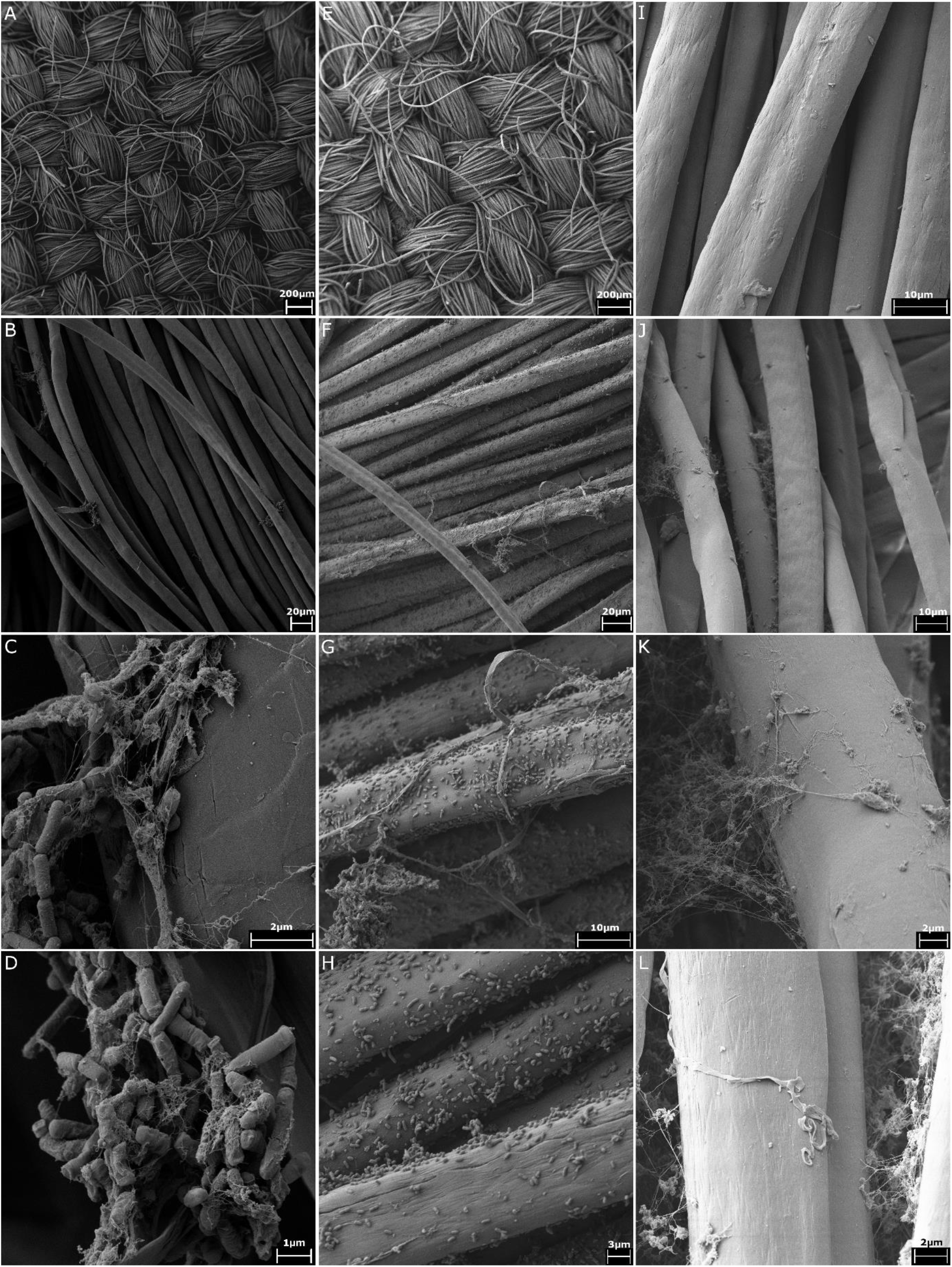
Scanning electron micrographs of NOMEX^®^ fabric which was co-incubated for six weeks with bacteria isolated from the ISS: A-D: Co-incubation with *Bacillus paralicheniformis*; E-H: Co-incubation with *Cupriavidus metallidurans*; I-L: Negative control of NOMEX^®^ fabric kept in sterile medium for six weeks. Precipitates found in the negative control were found to be remnants of the medium/particles and did not contain microbial cells.

*B. paralicheniformis* did neither adhere to the untreated nor to the eloxated aluminum alloy. However, the untreated aluminum alloy which was co-incubated with *B. paralicheniformis* showed sporadical signs of corrosion compared to the untreated negative control incubated in sterile medium. All eloxated aluminum surfaces showed attached debris regardless if the incubation in the sterile medium was performed under oxic or anoxic conditions. Single cells of *C. metallidurans* also attached to the surfaces of untreated and eloxated aluminum alloys and their co-incubated aluminum alloys had a unique background surface pattern, which was distinct from their respective negative controls. Under anoxic conditions, *C. avidum* formed a biofilm attached to the surface of both, untreated and eloxated, aluminum alloys.

## Discussion

In this study, we aimed to exploit the microbial information obtained from three sampling events aboard the International Space Station with respect to: i) microbial sources, diversity and distribution within the ISS, ii) functional capacity of microbiome and microbial isolates, iii) extremotolerance and antibiotics-resistance (compared to ground controls), and iv) microbial behavior towards ISS-relevant materials (biofilm formation, potential degradation or corrosion).

### Study limitations

Spaceflight experiments especially suffer from the limitations given by the circumstances that cannot be influenced by the scientists. This affected e.g. the number of samples to be taken (limited mass of payload), the sampling procedure (compatible to microgravity conditions and safety requirements), selection of sampling time points (according to assigned crew time and the overall mission planning), and the delivery duration of the samples to the laboratory. Being aware of these circumstances, experiments were planned accordingly (> 5 years implementation phase), and numerous controls were run to ensure the integrity of the information retrieved.

### Microbiome sources and context

Based on our observations and previous reports (10), we confirm a mostly human-associated microbiome aboard the ISS (62). Other proposed sources are cargo delivery (due to the detected overlap of cleanroom/ cargo and ISS microbial community), food (such as seasoning, dried fruits, nuts and herbs, or even probiotics (63) (as indicated by the presence of e.g. *Bacillus* and *Lactococcus* signatures), and potentially the personal belongings brought to the ISS (possibly reflected by the increased diversity in the personal sleeping unit microbiome). It shall be noted, that cargo deliveries are cleaned or even disinfected before upload (64,65), but an international standard for these procedures does not exist and thus the cleanliness of the cargo delivery might vary.

As a consequence, the ISS microbiome was characterized by a predominance of human (skin)-associated *Staphylococcus, Corynebacterium* and *Streptococcus* signatures (66). In general, these microorganisms are typical indicators for confined indoor environments (cleanrooms, space stations, hospital areas such as intensive care unit, operating rooms; (10)), emphasizing the substantial contribution of the human microbiome (see also (13)).

In more detail, all top 20 genera described in the hospital study (66) were also detected in the entire ISS microbiome (mostly also under top 20). *Enhydrobacter* (Pseudomonadales, a typical environmental species, (67)) was the only hospital top 20 genus which was not detected by the molecular approach in this study, but an *Enhydrobacter aerosaccus* isolate was obtained from the Columbus RGSH sample in session B.

Within the top 20 list of the ISS microbial signatures, *Haemophilus, Aerococcus, Stenotrophomonas, Gemella, Bacteroides, Actinomyces, Veillonella, Granulicatella, Blautia, Propionibacterium* and *Enterobacter* could not be found in the top 20 hospital list, indicating that those were relatively higher abundant in the ISS or even specific for this location.

All of these genera are typical human-associated microorganisms and thrive in the oral/respiratory tract (e.g. *Haemophilus*), on human skin (e.g. *Propionibacterium*) or the human gut (e.g. *Blautia)*. Some of these microorganisms have opportunistic pathogenic potential, as also pointed out for *Enterobacter* species isolated from the ISS WHC (68). We obtained in total eleven ISS isolates belonging to biosafety risk group S2 during this study, including *Pseudomonas putida* isolated from the RGSH in Node2 and isolates of the *Bacillus cereus/anthracis/thuringensis* clade retrieved from the RGSHs in Node2 and Columbus, from the hand grips in Columbus, and from the sleeping unit in Node2. However, most of these are typical human-associated bacteria which have only opportunistic pathogenic potential. Especially in the light of a weakened human immune system in space conditions, the presence and abundance of such opportunistic pathogens has of course to be carefully monitored, but as these do thrive in and on the human body and are shed into the environment by the crew itself, such opportunistic pathogens will always exist in built environments and their presence *per se* is not alarming (10).

### ISS microbiome diversity, biogeography and dynamics

The ISS microbiome was not found to be stable in composition and diversity, although a vast core microbiome existed over time and independent from location. However, specific microbial patterns could be identified for various functional areas within the station, as e.g. the WHC and RGSH showed the highest number of unique RSVs, and bacterial and in particular archaeal signatures were found to be specifically indicative for certain areas. This is in agreement with the observations of Ruiz-Calderon *et al*. that, increasing with the level of human interaction, indoor surfaces reflect space use and, besides, show increased content of human-associated microbial signatures (69).

Fluctuations in diversity and composition were detectable when two different time points (session A and B) were compared. It shall be mentioned, that no crew exchange took place during this period, but two cargo deliveries docked within this time frame (SpaceX and Soyuz), which could have influenced the microbiome composition. However, we detected an increase in specifically human (gut)-associated microorganisms (*Escherichia/Shigella, Lachnoclostridium* etc.) over the sampling period, of which the reason is unknown.

A different picture was obtained from the cultivable diversity of the ISS microbial community, with *Micrococcus yunnanensis, Bacillus hunanensis, B. megaterium* and *B. safensis* and *Staphylococcus epidermidis* being cultivable from both sampling sessions, representing hardy (spore-forming) and human skin-associated microorganisms, whereas typical gut-associated microorganisms could not be retrieved by our enrichments, as our cultivation approaches were designed to target rather environmental, extremotolerant microbes. Thus, the cultivation- and molecular-based microbial community analysis is not fully comparable due to the limitations in cultivation efforts.

Archaeal signatures were detected in 14 of the 24 ISS samples (universal and specific approach). Most earlier studies on the ISS microbiome ignored the possible presence of Archaea, but some of the more recent studies also reported the detection of Archaea aboard the ISS but did not further discuss their existence (19,21). We found, that archaeal signatures were nicely reflecting the frequency of human contact and the type of surface (see Fig. 5). The detected sequences of Thaumarchaeota, Woesearchaeota and *Methanobrevibacter* (Euryarchaeota) have all been attributed to the human microbiome before (70–72).

### Functional capacity of microbiome and microbial isolates, extremo-tolerance and antibiotics-resistance (compared to ground controls)

In our study, we confirmed the pangenome-based observation of Blaustein *et al*. (24), on genomic, but also on isolate and resistance-pattern level that ISS microorganisms are not necessarily more extremophilic or antibiotic resistant than their ground-relatives with respect to growth behavior, or antibiotics resistances, nor their genomic inventory.

We rather argue that the ISS environment supports selection of the best-adapted microorganisms (e.g. spore-formers) towards the partially extreme physical and chemical environmental conditions (e.g. radiation, alkaline cleaning agents), but does not induce general changes in the physiological nor genomic capacities of microbes. Thus, were not able to confirm the null hypothesis that strains obtained from the ISS are more extremotolereant/extremophilic than closely related strains from Earth regarding the upper and lower boundaries of their temperature and pH growth ranges. With the exception of *Bradyrhizobium viridifuturi* pH5_R2_1_I_B, all tested ISS isolates were able to grow at pH 9 or higher, which might be due to a selection pressure caused by alkaline cleaning reagents used on board the ISS.

Moreover, the data presented here show that the molecular detection of antibiotic resistance genes, while being a good approximation of the resistance potential of an organism or microbial community, does on the one hand overestimate the antibiotic resistance potential (as some resistance genes might not be expressed at all). On the other hand it does not necessarily cover all antibiotic-resistances which a microorganism actually has. Thus we advise coupling traditional cultivation approaches with molecular investigations to retrieve a full picture of the situation.

### Microbial surface interaction with ISS materials

Although we could not confirm an increased threat of ISS microbiome towards crew’s health, we observed that surface interaction is critical for the microbial community aboard. A variety of ISS surface materials are composed of metals, including alloy EN AW 2219 (aluminum copper magnesium), which might stress interaction with metal ions and settlement. In our co-incubation experiments, we could confirm that ISS microbial isolates can adhere and grow on metal and in particular textile surfaces (NOMEX fabric), where local moisture (e.g. condensate) could support biofouling, biofilm formation and material damage through acid production (6,73–75). This is particularly important with respect to fungal growth, as these might affect human health indirectly by causing allergic reactions and asthmatic responses (17,76–78).

### Conclusion: Microbes aboard: Reason for concern?

Although we cannot fully exclude a threat of the ISS microbiome towards crew health (in particular in interaction with the weakened human immune system) our data do not indicate direct reason for concern. However, we raise special attention to the microbial-surface interaction problem in order to avoid biofouling and biofilm formation, which could directly impact material integrity and indirectly human health and therefore pose a potential risk to mission success.

## Acknowledgements

We are thankful for financial support by the FFG (Austrian Research Promotion Agency, Project No. 847977) and the European Space Agency (ESA) for financing and realization of this space project as part of the ELIPS program (ILSRA-2009-1053 ARBEX). We are very grateful for the scientific and technical support by Lobke Zuijderduijn, Stefanie Raffestin (sampling in Kourou), the BIOTESC team (Lucerne University of Applied Sciences), the ISS crew and all other involved members of space agencies and attached institutes. We thank Elisabeth Grohmann for providing a *Staphylococcus* strain isolated from the Concordia station. PhD student Maximilian Mora was supported by the local PhD program MolMed.

